# Maternal infection causes dysfunctional BCR signaling in male offspring due to aberrant Xist expression

**DOI:** 10.1101/2023.02.13.528357

**Authors:** Lisa C. Gibbs, Juan M. Oviedo, Bartholomew N. Ondigo, Keke C. Fairfax

## Abstract

Infections during pregnancy with pathogens such as helminths correlate with altered immune responses to common childhood immunizations. However, the molecular mechanisms that underlie this remain unknown. Using our murine model of maternal schistosomiasis, when immunized, males from infected mothers had a lower frequency of antigen-specific germinal center B cells and downregulation of transcripts downstream of BCR signaling compared to males from uninfected mothers. This is driven by a reduction in developing B cell populations within the bone marrow of pups from infected mothers. Males from infected mothers were impacted to a greater extent than their female littermate counterparts. We found this defect to be caused by aberrant expression of the long non-coding RNA *Xist* in males leading to dysregulated Igα expression on developing B cells. This, for the first time, links dysfunctional BCR signaling with *Xist* expression, while also proposing a detrimental function for *Xist* expression in males.

**One sentence summary:** *Xist* expression in males decreases BCR reactivity.

## INTRODUCTION

Despite high vaccination coverage, infectious diseases remain prevalent around the globe, with vaccine efficacy differing by regions (*1, 2*). Specifically, in some regions of Africa where parasitic infections are endemic, efficacy to common childhood vaccines is lower than in regions where parasitic infections are not as prevalent (*3, 4*), raising the possibility that this is caused by bystander immunosuppression driven by parasite infection (*5, 6*).

Schistosomiasis is caused by a parasitic helminth that induces Th2 skewing during chronic infection. While an inappropriate excess of Th2 cytokines can prolong virus clearance (*7*), these cytokines can also decrease chronic, autoimmune-induced inflammation (*8*). Maternal parasitic infections can mirror these disease outcomes in offspring, such as decreased responsiveness to antigens and protection from the development of some allergies and other chronic diseases (*9-12*). Yet, a decrease in Th2 cytokines is observed in mouse models of maternal schistosomiasis, due to epigenetic regulation of the transcription of these cytokines (*10, 13*). This suggests the immunosuppressive responses seen in offspring born to infected mothers is not due to excessive offspring Th2 cytokine production, but other, undefined mechanisms.

Maternal helminth infections, specifically schistosomiasis, can have an impact on offspring vaccine efficacy to common childhood vaccines (*11, 12, 14*). Maternal schistosomiasis has a prevalence rate varying between 1-60% across regions of Africa (*15*). In these studies, the children tested have not been infected with schistosomiasis, yet they have decreased vaccine efficacy (*14*), increasing the risk of a vaccine preventable disease outbreak in these communities. We have previously published using a murine model of maternal schistosomiasis that pups from schistosome infected dams have transcriptional dysregulations of key germinal center genes after vaccination, such as *Ebf1* and *Junb* (*13*), restricting the germinal center reaction and decreasing vaccine titers.

In this study we use this established murine model of maternal schistosomiasis (*13*) to mechanistically understand how maternal infections impact offspring immunity. First, we found that males born to *Schistosoma mansoni* infected mothers, and not females, have decreased responsiveness to vaccination, and validated this finding with comparable, human immno-epidemiological data. This deficiency was due to a defect in steady state bone marrow B cell development. Developing B cells from males from infected mothers have lower surface expression of Igα, a transmembrane protein necessary for BCR signaling. We found that low Igα expression in males was being caused by aberrant *Xist* expression. *Xist* is a long non-coding RNA that canonically coats one X chromosome in females for dosage compensation. Here, we show that aberrant *Xist* expression in males, caused by maternal schistosomiasis, decreased Igα on B cells and leads to decreased responsiveness to antigens. This study is the first to provide a molecular mechanistic understanding explaining decreased vaccine efficacy during maternal schistosome infection, while also defining a function of *Xist* in males.

## RESULTS

### Males from *S. mansoni* infected mothers have lower tetanus vaccine efficacy

To understand the mechanism behind lower vaccine efficacy seen in children born to schistosome infected mothers (*14*), we employed our previously published maternal *Schistosoma mansoni* infection murine model (*13*). 4get homozygous females, (GFP IL-4 reporter mice), were either sham-infected or infected with a low dose of *S. mansoni*. After the start of the egg laying at 6 weeks, females were paired to KN2 homozygous males, which produce human CD2, a transmembrane reporter protein, instead of IL-4, resulting in dual IL-4 reporter pups (*16*). At 28-35 days old, aged-matched pups from sham-infected (uninfected or control) and schistosome infected dams were immunized with a commercial tetanus/diphtheria (Td) vaccine (Fig. 1A). We have shown that mice born to infected mothers have lower frequencies of total germinal center (Fig. S1A) (GC) B cells, memory B cells, and plasma cells (*13*). To examine the antigen specific B cell response, we used fluorescently conjugated tetanus toxoid (Fig. S1B). Both males and females from infected mothers have decreases in tetanus-specific B cells within the GC (Fig. 1B), complementing the lower bulk B cell GC frequency. We found an accumulation of phospho-STAT3 in males from infected mothers, but not in females (Fig. 1C). This accumulation is commonly seen in pathogenic, atypical GC formations, such as during lupus and large B-cell lymphoma (*17, 18*), suggesting an abnormal GC response.

**Figure 1:**
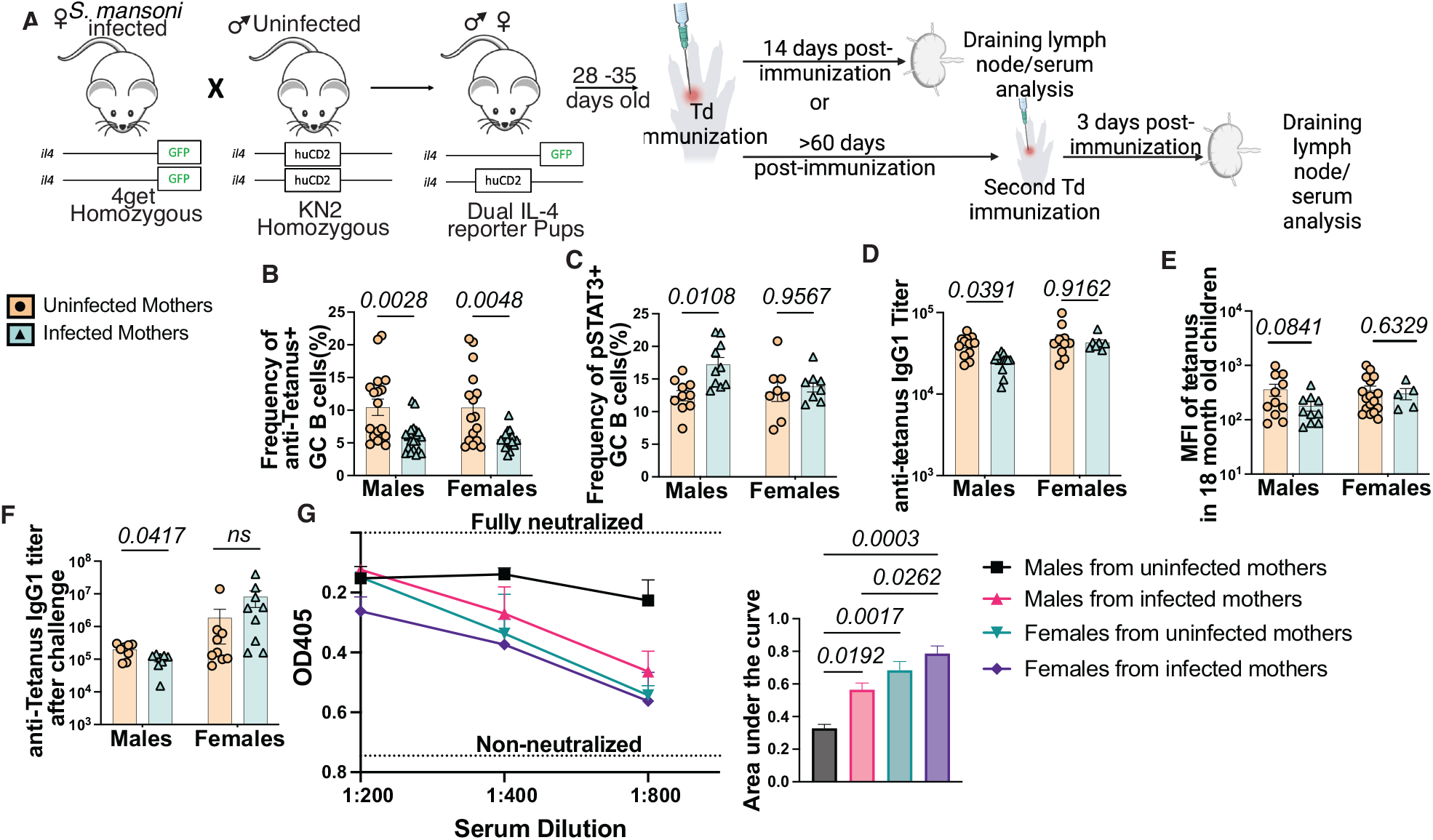
Sex-specific defects in GC response after maternal *S. mansoni* infection leads to lower vaccine efficacy in males. (**A**) Model schematic for generation of pups born to *Schistosoma mansoni* infected mothers. Fourteen days post vaccination, or after rechallenging, draining lymph nodes and serum were assessed. (**B**) Flow cytometry of anti-tetanus positive GC B cells (CD19^+^GL7^+^CD95^+^) in draining lymph node of 14 days post subcutaneous Td immunization. Each point represents a mouse. Experiments performed 3 times and represents >5 litters. (**C**) Flow cytometric frequencies of phospho-STAT3 positive GC B cells. Experiment representative of >4 litters. (**D**) Anti-tetanus IgG1 titers calculated from ELISA from serum of pups from control and *S. mansoni* infected mothers 14 days post Td vaccination. (**E**) Data analyzed from data set previously collected from antibody titers from serum of children in Kenya born to schistosome infected and control mothers, using published cut off values. (**F**) Anti-tetanus IgG1 titers calculated from ELISA from serum of pups from control and *S. mansoni* infected mothers 3 days post-secondary Td vaccination. (**G**) Tetanus neutralizing assay using serum from Figure 1F. All statistics determined by two-way ANOVA.

Males from infected dams also have decreased anti-tetanus IgG1 titers in serum, measured by ELISA (Fig. 1D) with no decrease in antibody avidity (Fig. S1C). Interestingly, we found this to also be true in children born to schistosome infected mothers. In western Kenya, male children from *S. mansoni* infected mothers have a trending decrease of anti-tetanus IgG1 titers (p= 0.0841), while titers of female children are not affected (Fig. 1E)(*14*), suggesting a sex-specific effect for tetanus responses in this population.

The function of immunization is to prime a host to mount a rapid response to homologous antigenic challenge, often characterized by high neutralizing antibodies. To test the recall ability of B cell pool, pups from control and *S. mansoni* infected dams were immunized with Td. Sixty days post primary immunization, mice were boosted with Td and response measured three days later. As expected, males from infected mothers had lower anti-tetanus IgG1 titers in serum after re-challenge compared to controls (Fig. 1F). When testing the protective nature of these antibodies by a tetanus neutralization assay, males from control dams had the highest neutralization rate. In comparison, males from infected mothers had significantly less neutralizing ability (Fig. 1G). Moreover, females from control and infected mothers had the least neutralizing potential, which is often observed humans with other vaccinations (*19, 20*).

Together these data show males from *S. mansoni* infected mothers have decreased vaccine efficacy to Td.

### B cell development during maternal *S. mansoni* infection is impaired after BCR recombination

To understand the origin of decreased GC B cells, we first wanted to determine if this was solely due to the GC reaction or if it is a systemic B cell defect that begins during B cell development. We found a positive correlation between the frequency of transitional and mature (T1) B cells to the frequency of naïve B cells in the periphery (Fig. 2A). As expected, developing B cells in the bone marrow have an impact of the number of mature B cells in the periphery.

**Figure 2:**
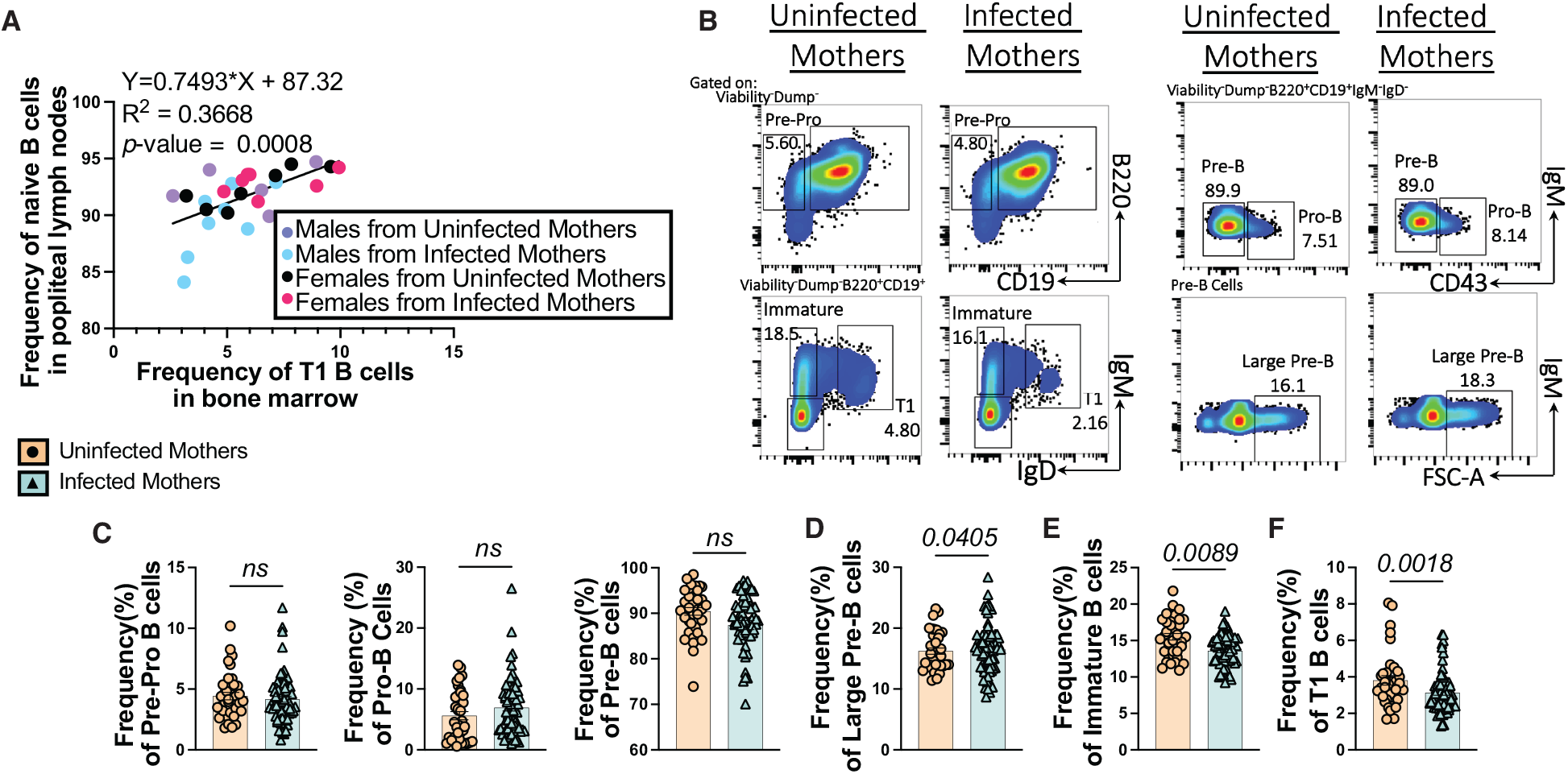
Pups from *S. mansoni* infected mothers have reduced developing B cell populations, causing less B cells in the periphery. (**A**) Correlation plot of the frequency of T1 B cells in the bone marrow compared to the frequency of CD19^+^IgM^+^IgD^+^ B cells in popliteal lymph nodes. Each dot represents a single mouse. Experiment performed two separate times, representative of four separate litters. (**B**) Concatenated flow cytometry plots showing gating strategy for bone marrow B cell populations. (**C**) Frequencies of pre-pro (left), pro (mid), and pre (right) B cells by flow cytometry from bone marrow of pups from infected and control mothers. Each dot/triangle represents a single mouse. Flow cytometry experiments represented by bar graphs performed >10 separate times, representative of >10 separate litters. (**D**) Frequency of large pre-B cells by flow cytometry from bone marrow of pups from infected and control mothers. (**E**) Frequency of immature B cells by flow cytometry from bone marrow of pups from infected and control mothers. (**F**) Frequency of transitional/mature (T1) B cells by flow cytometry from bone marrow of pups from infected and control mothers. All statistics determined by student’s t-test.

To begin surveying the developmental B cell populations to determine if there was a defect, we used the following flow cytometric gating strategies for various stages of B cell development on cells extracted from femurs of pups (Fig. 2B) (*21-23*). There was no change in whole bone marrow cellularity (Fig. S2A) or frequency of pre-pro B cells, pro-B cells, or pre-B cells between pups from infected and control mothers (Fig. 2C). In contrast, there is an increase in large pre-B cells (Fig. 2D), the proliferative stage of pre-B cell development (*24*). These cells also have successfully rearranged Igμ of the Ig heavy chain, while still expressing the surrogate light chain (SLC) (*25*), forming the pre-BCR. While the pre-BCR and pre-B cell stage is an important checkpoint for B cell lymphopoiesis (*26*), a cellular increase is not conserved throughout the rest of B cell development in pups from infected mothers. In fact, pups from schistosome infected mothers have decreases in immature (Fig. 2E) and transitional/naïve mature (T1) (Fig. 2F) B cells, resulting in a deficiency of about 100,000 immature and T1 B cells per femur (Fig. S2A). Additionally, this defect in immature and T1 B cells is not IL-4 dependent, as IL-4 deficient pups born to IL-4 competent schistosome infected mothers still have decreases in these population (Fig. S2B). Importantly, this defect does not seem to be genotype dependent, as mixed background pups born to schistosome infected mothers on a C57BL6/J background and fathers on a BALB/cJ background also share this defect (Fig. S2C).

This suggests that a large percentage of large pre-B cells fail to develop into immature B cells, and subsequently do not become T1 or mature B cells due to failed checkpoint passage.

Together, these data show that pups born to schistosome infected mothers have a decrease in immature and T1 B cells in the bone marrow, caused by failure to pass the SLC checkpoint during the pre-B cell stage, resulting in a decreased pool of B cells within the periphery.

### Sex-specific B cell signaling defects begin in pre-BCR signaling during maternal schistosomiasis

Males from schistosome infected mothers cluster together towards the bottom of the correlation plot when comparing developmental, bone marrow B cells to peripheral B cell populations, often falling below the best-fit line (Fig. 2A). This suggests that males from infected mothers are more impacted than female littermates. We have previously published that EBF1, a transcription factor necessary for B cell proliferation, development and function (*27*), is reduced by both transcript and protein levels in naïve, peripheral, and GC B cells (*13*). Specifically for pre-B cells, EBF1 directly regulates the transcription of the proteins that encompass the SLC of the pre-BCR, λ5 and V_preB_ (*28, 29*). Indeed, there is a decrease in EBF1 expression on pre-B cells (Fig. 3A), coupled with an expected decrease in V_preB_ expression (Fig. 3B), while females from infected mothers are unimpacted compared to females from control mothers. Males from infected mothers also have decreased *Id3* transcription (Fig. 3C). EBF1 expression is necessary to downregulate *Id3* transcription before continuing through B cell development (*30*), suggesting that during maternal schistosomiasis, males and not females from infected mothers have impaired pre-BCR signaling cascades, leading to complications in B cell differentiation.

**Figure 3:**
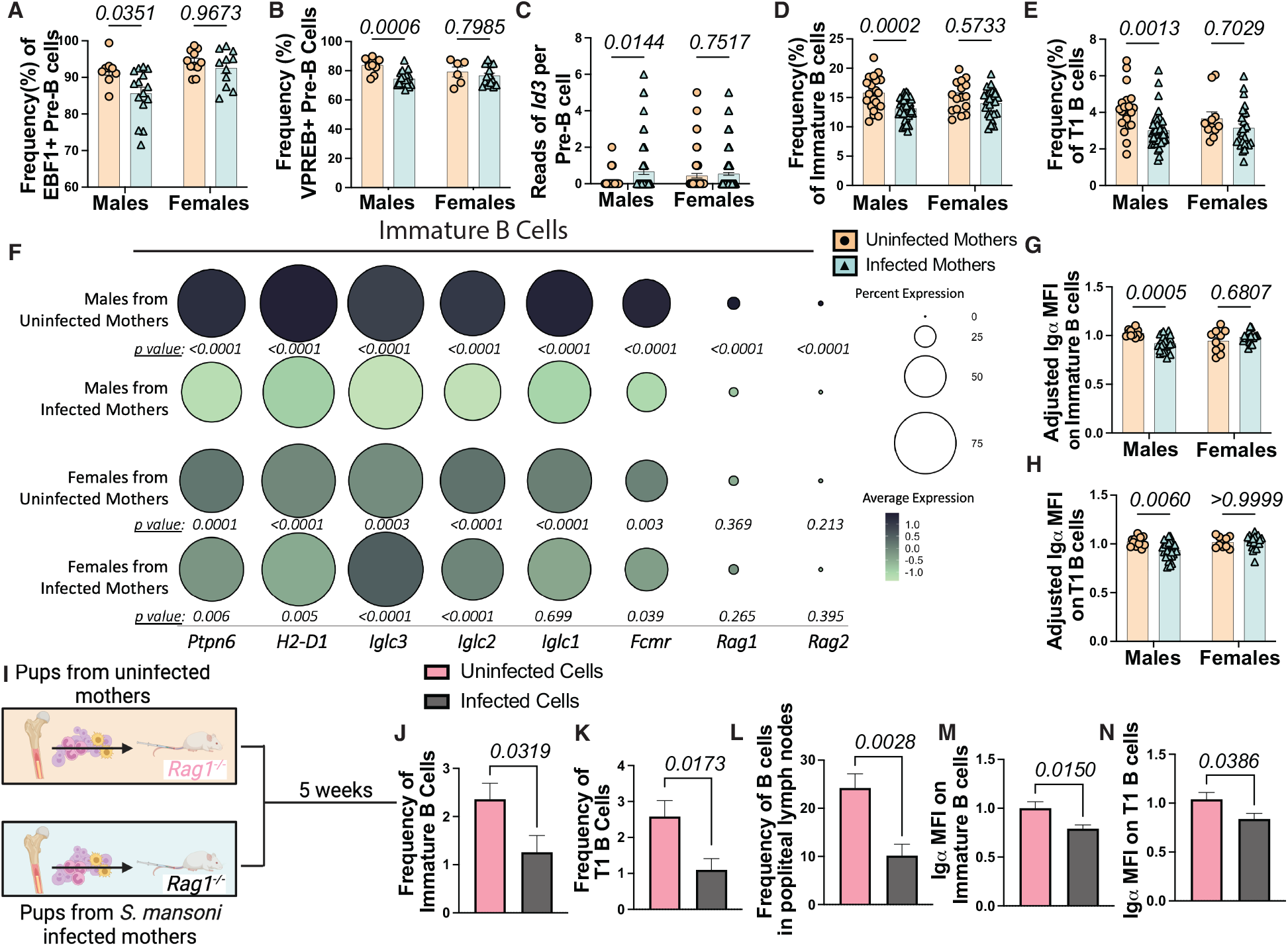
Males from *S. mansoni* infected mothers have lower Igα surface expression on bone marrow B cells. (**A**) Frequencies of EBF1^+^ and (**B**) V_preB_^+^ pre-B cells from pups from *S. mansoni* infected and control mothers by flow cytometry. Each dot represents a single mouse. Experiment performed >3 separate times, representative of >5 separate litters. Statistics calculated by two-way ANOVA. (**C**) Number of reads of *Id3* transcript per pre-B cell from single cell RNA sequencing of mature myeloid cell depleted bone marrow. Pre-B cell cluster subset before calculating differentially expressed genes and *p*-values. Each group represents at least 3 mice. (**D**) Flow cytometric frequencies of immature and (**E**) T1 B cells from bone marrow of age matched pups from *S. mansoni* and control mothers from Fig. 2, split by sex. Statistics calculated by two-way ANOVA. (**F**) DotPlot from single cell RNA sequencing of mature myeloid cell depleted bone marrow. Immature B cell cluster subset before calculating differentially expressed genes and *p*-values. *p*-value shown is for comparison to group above compared to males from infected dams. Each group represents at least 3 mice. Transcripts listed below. Average expression, or average transcript expression per cell, is depicted on a color scale with black being highest expression and green being decreased expression. Dot size correlates with percent expression, or number of cells expressing each transcript. (**G**) Mean fluorescent intensity (MFI) of Igα on immature and (**H**) T1 B cells by flow cytometry. Experiments performed >4 times, representative of >6 litters. Statistics determined by two-way ANOVA. (**I**) Model schematic for adoptive cell transfer. (**J**) Frequency of immature and (**K)** T1 B cells in the bone marrow of *Rag1*^*-/-*^ mice after receiving bone marrow from 4getKN2 males from control and infected mothers. Experiments performed 3 times, representing >6 litters. (**L**) Frequency of CD19^+^ B cells in lymph nodes of *Rag1*^*-/-*^ recipients. (**M**) Adjusted MFI of Igα on immature and (**N**) T1 B cells of *Rag1*^*-/-*^ recipient. Statistics determined by student’s t-test.

### Males from *S. mansoni* infected mothers have lower Igα on the surface of developing B cells

To determine the impact of defective development after V(D)J recombination in males from infected mothers, developing B cell populations downstream of pre-B cells were examined. Immature B cells, which arise after Igκ recombination, have about a one-third decrease in males from infected mothers compared to males from control mothers, which is not seen in females (Fig. 3D). Consequently, males from infected mothers, but not female, have lower T1 B cells (Fig. 3E). Interestingly, this drop in T1 B cells that develop from immature B cells is a loss of almost half, suggesting a more stringent negative selection process from an immature B cell to T1 B cell (*31*).

To look at BCR selection differences, single cell RNA sequencing (scRNASeq) was performed on developing B cells. Briefly, bone marrow was harvested from pups from control and infected mothers. CD3^+^CD4^+^CD11b^+^CD11c^+^ TER119^+^ cells, termed myeloid-depleted bone marrow for simplicity, were excluded by FACS sorting, while developing and mature B cells were sequenced (Fig. S3). The immature B cell cluster was subsetted and expression level of genes important for selection and receptor editing, such as *Ptpn6* (*32*), *Fcmr* (*33*), *Rag1* and *Rag2* (*34*), along with genes comprising the light chain of the BCR (*Iglc3, Iglc2, Iglc1*) and MHCII (*H2-D1*) were examined. Unsurprisingly, males from infected mothers downregulated these genes to a greater extent than in females from infected mothers (Fig. 3F), suggesting a skewing away from conventional receptor editing, or a secondary chance at recombination (*35*).

Differences in selection can arise from defects in signaling, either caused by the signaling components themselves, or by unfavorable V(D)J recombination. Both processes lead to an increase of B cells being selected out of the pool that egress from the bone marrow and limit the number that get to the periphery and can respond to antigen. We explored both possibilities to determine where the selection defect comes from. Igα and Igβ, two transmembrane proteins, form the BCR complex with IgM on immature B cells and are important docking sites for signal transduction (*34, 36*). We looked at surface expression of Igα, which is indispensable for the induction of the BCR signaling cascade (*37*). As expected, males from infected mothers have decreased mean fluorescent intensity (MFI) of Igα on the surface of immature (10.3% reduction, Fig. 3G) and T1 (8.3% reduction, Fig. 3H) B cells, coupled with a reciprocal increase of immature and T1 B cell with low surface expression of Igα (Fig. S4A, S4B).

To determine if these defects were cell intrinsic or being caused by the bone marrow microenvironment, bone marrow cells from males from uninfected and infected mothers were transferred to *Rag1*^-/-^ mice that do not have mature B or T cells (Fig. 3I). Mice that received bone marrow from males from infected mothers had a decrease in immature (Fig. 3J), T1 (Fig. 3K), and peripheral (Fig. 3L) B cells. Additionally, the decrease of Igα on immature (Fig. 3M) and T1 (Fig. 3N) was retained. This demonstrates that the B cell defects seen in males from infected mothers is cell intrinsic, suggesting a long-lived transcriptional or epigenetic mechanism controlling this dysfunction.

Lower Igα and subsequent B cell developmental populations indicate a defect within the BCR that restricts signaling, mirroring a phenotype where heavy chain expression is mutated (*38*) and B cells become non-functioning and unable to activate. This suggests that Igα low B cells are less fit for survival, leaving the males from infected mothers with a decreased pool of mature B cells, as seen in the periphery (Fig. 2A).

### Males from infected mothers have an increase in unfavorable clonotypes

Next, we asked if these B cells were less fit solely due to defective Igα signaling, or if there was also a defect in V(D)J recombination. To test the outcomes of recombination, clonotypes of immature and T1 B cells were examined by single cell V(D)J sequencing (Fig. S5A, S5B).

Although Igα functionality does not affect the ability to undergo V(D)J recombination (*39*), when comparing clonotypes of males from infected mothers to males from uninfected mothers, males from infected mothers have an expansion of unique clones, shown on the Y-axis (Fig. 4A). These unique clonotypes are not due to the pro-inflammatory environment during maternal infection because when comparing littermate females from infected mothers to females from uninfected mothers, most of their clonotypes are shared (Fig. 4B). Additionally, when comparing females and males from infected mothers, there is an increase in unique clonotypes in the males (Fig. 4C), again suggesting that these are not due solely to maternal factors and are influenced by the sex of the pup.

**Figure 4:**
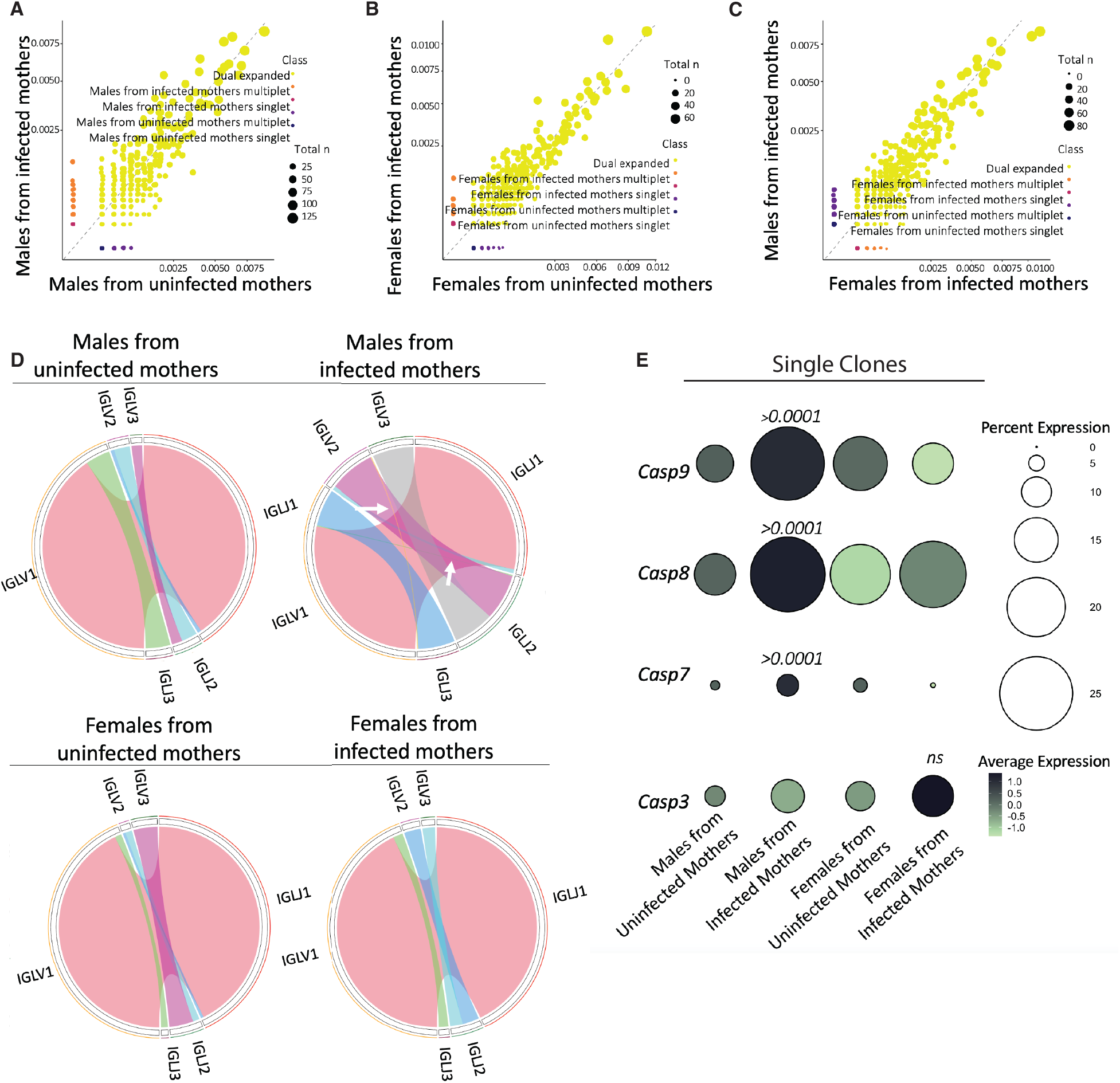
Defects on B cells of males from infected mothers lead to greater unfavorable clonotypes. (**A**) ScatterPlots generated by scRepertoire comparing clonotypes of immature B cells between males from *S. mansoni* and control mothers and (**B**) females from *S. mansoni* and control mothers and (**C**) males and females from infected mothers. Single cell V(D)J sequencing was performed on B220^+^CD19^+^IgM^+^IgD^+/-^ bone marrow from pups from *S. mansoni* infected and control mothers. Each group represents at least 3 mice. Class groups clonotype based on if they are single or expanded and for which sample. Total n shows how many sequenced BCRs are represented by each dot, corresponding to dot size. (**D**) Gene usage charts from VDJView from each sample group from single cell V(D)J sequencing showing λ gene pairings between variable and joining regions. White arrows on males from infected mothers indicate pairings not present in other samples. (**E**) Single clones were subset before calculating differentially expressed genes and *p*-values. Transcripts analyzed are listed to the left of the dots. *p-*values listed represent *p*-value between males from infected mothers compared to other three groups. Average expression and percent expression are the same as explained above.

Because of the light chain transcriptional defects, we identified in Fig. 3F, we looked at light chain gene usage. While males from uninfected mothers and females from infected and uninfected mothers had one main pairing (*Iglv1* to *Iglj1*), there were also three secondary pairings that occur (Fig. 4D). Males from infected mothers uniquely have the increased ability to pair *Iglv2* to *Iglj3* and *Iglv1* to *Iglj2*, both of which are not observed in the other groups (Fig. 4D). This suggests that these pairings, because of their absence or rarity in other groups, are not highly occurring during normal B cell devolvement. This could mean they are either non-functional or autoreactive, and these possibilities will be examined in the future.

These unique clonotypes are not optimal because of the high expression of *Casp9, Casp8*, and *Casp7* in these cells (Fig. 4E). After unsuccessful BCR crosslinking, such as after failed selection, cell death can be mediated by the Caspase-8 pathway (*40*). This induces the formation of the apoptosome, which results in the activation of Caspases 3,7, and 9 (*41, 42*), effectively inducing apoptosis. Non-expanded clones from male pups born to infected mothers had significantly higher expression of *Casp8, Casp9*, and *Casp7* compared to non-expanded clones from other groups, indicating a higher rate of cell death after BCR crosslinking. Overall, males from infected mothers have differential light chain pairings, leading to an increase of unique clonotypes and a higher rate of cell death.

### Males from infected mothers have increased *Xist* expression

There are well-known sex biases in the immune response thought to be due to X-linked genes and/or sex hormones (*43*). Because sex hormone levels are low until about 40 days of age after birth (*44*), the role of sex hormones was not considered in this study. Focusing on X-linked genes, we found an increase in the expression of the long non-coding RNA *Xist* in bone marrow B cells in males from infected mothers (Fig. 5A). *Xist* is known to regulate dosage compensation in females by coating one X chromosome. Also, in females, *Xist* plays a pivotal role in B cell activation (*45*) and isotype switching (*46*). To determine if *Xist* expression begins before B cell maturation, we sorted common lymphoid progenitors, or the earliest lymphoid committed cell stage, and visualized *Xist* using RNA FISH (fluorescence in situ hybridization) (Fig. 5B). Males from infected mothers have a higher percentage of *Xist* positive cells (Fig. 5B) as compared to age-matched males from uninfected mothers. Female mice from infected mothers have similar *Xist* expression as controls. This shows that *Xist* expression is induced in males from infected dams from at least the common lymphoid progenitor cell level and is being maintained. *Xist* is normally transcribed in *cis* with its antisense, *Tsix*, which is essential for the degradation of both (*47*). Normally, there is increased expression of *Tsix* in comparison to *Xist* because it can persist on the active X chromosome (*48*). This is what is observed in males from uninfected mothers and females from infected and uninfected mothers on a per cell basis, but males from infected mothers have an almost equal ratio of each (Fig. 5C). Increases in *Tsix* levels usually acts as a repressor for *Xist* accumulation (*49, 50*), so a change in ratio as seen in males from infected mothers allows for *Xist* accumulation and aberrant *Xist* binding.

**Figure 5:**
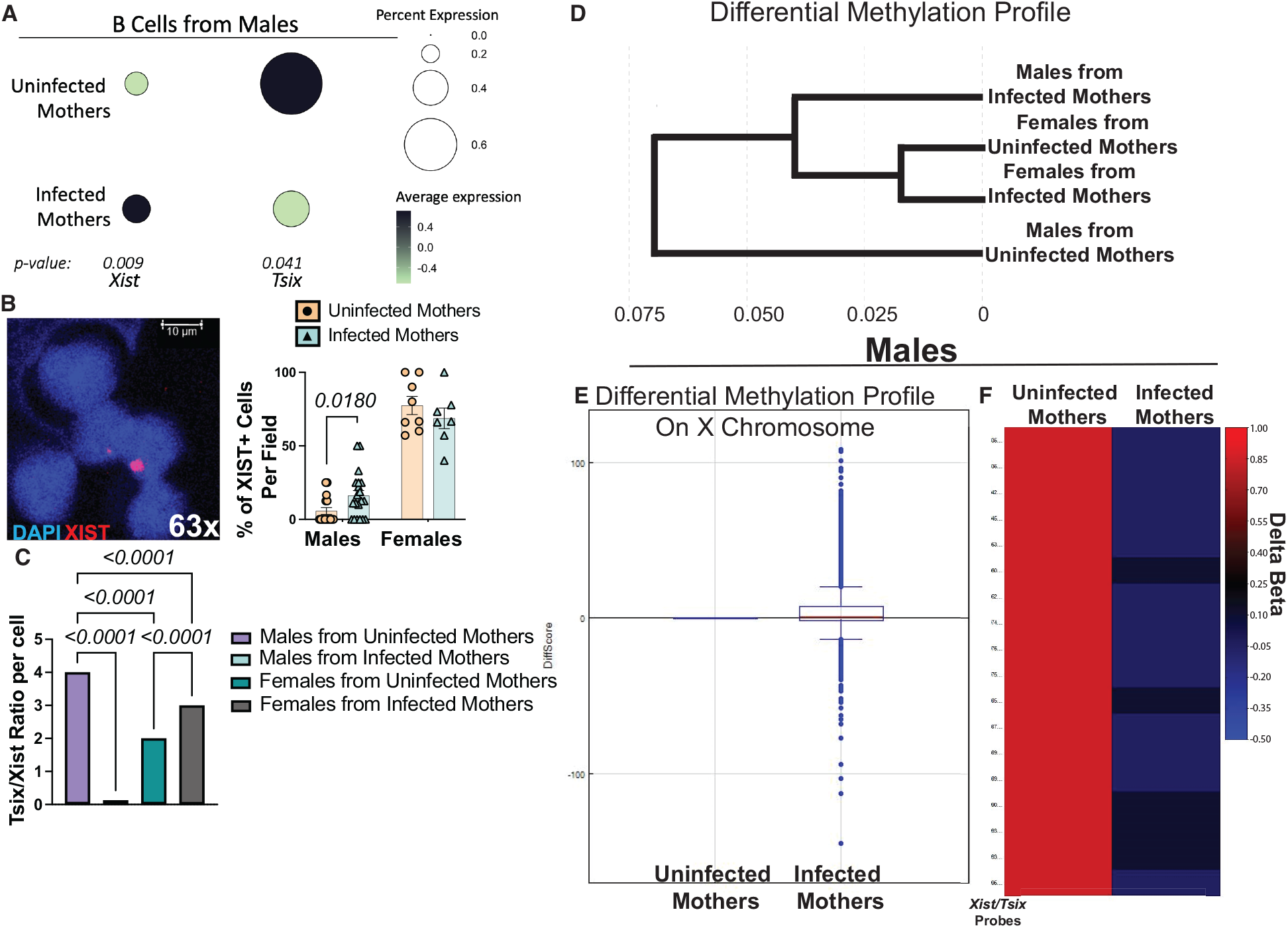
Differential methylation on the X chromosome allows for increased *Xist* expression in males from *S. mansoni* infected mothers. (**A**) DotPlot from data generated by scRNAseq showing *Xist* and *Tsix* from bone marrow B cells from males from *S. mansoni* infected and control mothers. B cells were subset by *Cd19* and *Ighm* positive expression. Each group represents at least 3 mice. Transcripts listed below. Average expression and dot size are as described above. (**B**) RNA FISH using fluorescently conjugate probes for *Xist*. Common lymphoid progenitors from bone marrow of male and female pups from *S. mansoni* and control mothers were sorted before staining. Images were taken at 63X with blue representing DAPI and red representing *Xist*. Left panel represents one field used to calculate number of positive cells (1 positive cell per 5 cells = 20%). Right panel shows percentage of positive cells with each point representing a single field. Statistics calculated by two-way ANOVA. (**C**) Ratio of *Tsix* to *Xist* per cell. B cells subset from scRANseq were evaluated for number of *Tsix* and *Xist* reads on a per cell basis. Number of *Tsix* reads were then divided by number of *Xist* reads per cell. Statistics calculated by one-way ANOVA. (**D**) Dendrogram created by Illumina™ Genome Studio after ChIRP using biotinylated *Xist* probes to pull down DNA interacting with *Xist*. DNA was extracted and then used for Illumina® Infinium HD Mouse Methylation Assay. Each group represents a pool of 4 mice. (**E**) Differential methylation analysis done in Illumina™ Genome Studio after ChIRP between males from *S. mansoni* infected and control mothers normalized to males from uninfected mothers. Each dot represents a probe. Diffscore is defined as differential methylation score. (**F**) Heatmap generated by Illumina™ Genome Studio showing delta beta (change in ratio of intensities between methylated and unmethylated) of all *Xist*/*Tsix* probes normalized to males from uninfected mothers.

To determine the role of *Xist* in males, whole bone marrow was subject to chromatin isolation by RNA purification (ChIRP) using probes for *Xist*. DNA was extracted from the *Xist* bound material and used for a methylation array. Comparing the methylation profiles of DNA bound to *Xist* of the whole genome, males from infected mothers cluster closer to females from control and infected mothers, than to males from control mothers (Fig. 5D). Isolating the X chromosome and normalizing to females from uninfected mothers, males from infected mothers have a spread of methylation comparable to males from uninfected mothers, but the DiffScore average is higher than males from uninfected mothers, indicating some differences between male groups Fig. 5E). Comparing probes specific for *Xist*/*Tsix*, males from infected mothers have a lower delta beta or ratio of unmethylated to methylated probes (Fig. 5F). Collectively, these data show that the regulation of *Xist* in male mice from infected mothers is unique in both its ability to escape degradation and to bind to differential positions along the X chromosome.

### Elevated *Xist* levels impair BCR expression

To functionally phenotype B cells with elevated *Xist* expression, *Xist* positive B cells from males from infected mothers were subsetted using the scRNASeq of myeloid depleted bone marrow. *Xist* positive cells have a transcriptional decrease in *Cd79a* and *Cd79b*, encoding for Igα and Igβ, respectfully (Fig. 6A). To validate this, high Igα expressing cells and low Igα expressing cells were sorted from the bone marrow of pups from infected mothers and RT-qPCR for *Xist* was performed. Interestingly, in males from infected mothers, the Igα low cells had an increase in *Xist* expression compared to the Igα high, or normal BCR expressing, cells (Fig. 6B). Notably, the females from uninfected mothers had varying *Xist* expression between the two B cell types (Fig. 6B), showing no correlation between Igα expression and *Xist*. Together, this suggests aberrant *Xist* expression, such as seen in males from infected mothers, causes B cells to have low surface Igα, and therefore less BCR signaling capacity.

**Figure 6:**
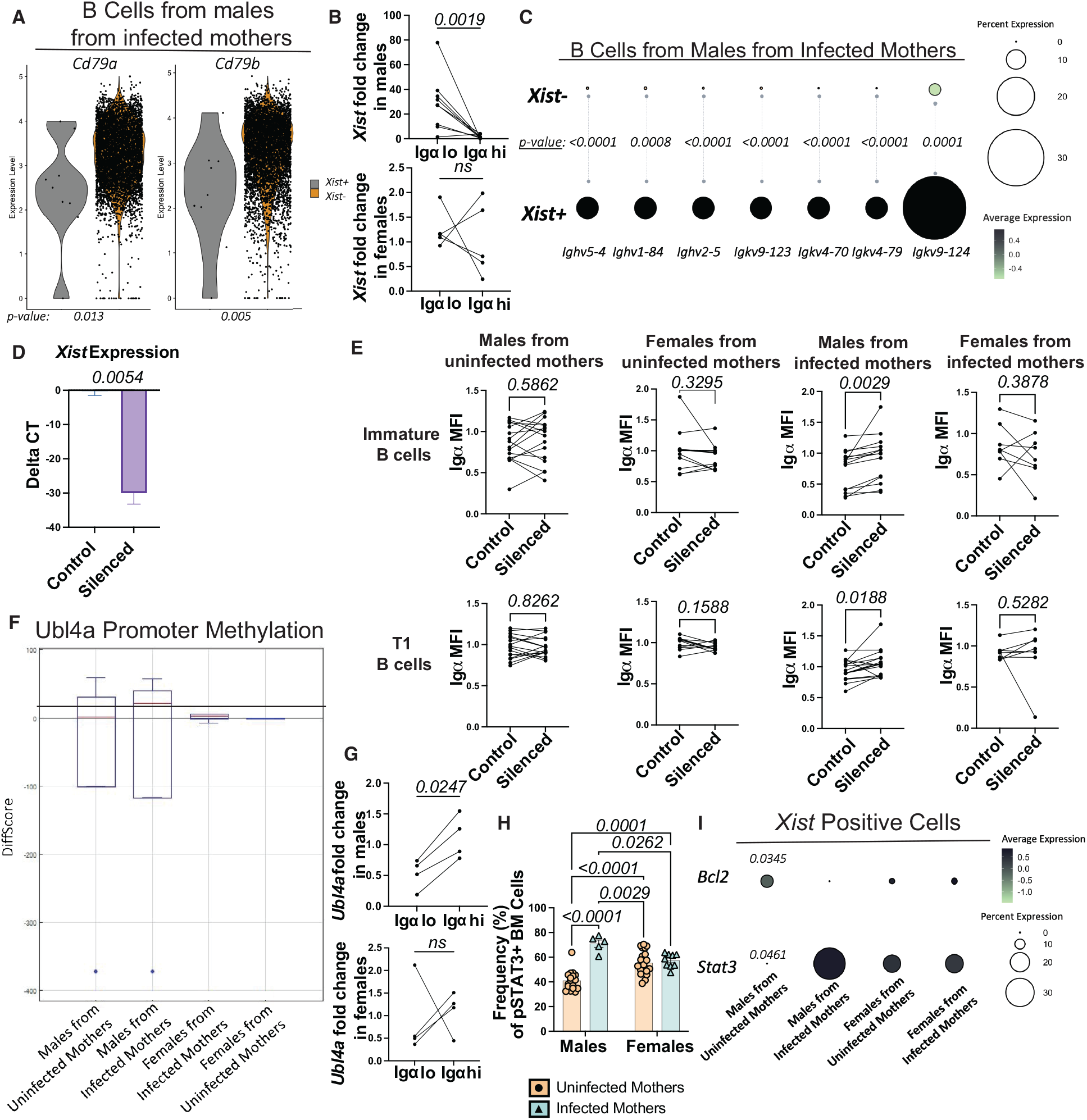
Elevated Xist expression in males from *S. mansoni* infected mothers causes decreased Igα expression. (**A**) Violin Plots showing *Cd79a* (Igα)(left) and *Cd79b* (Igβ)(right) expression level on B cells from males from *S. mansoni* infected mothers subset from bone marrow B cell scRNAseq. Each dot represents a cell. *p-*values are listed below. (**B**) Fold change of *Xist* on bone marrow B cell populations sorted by Igα high and low surface expression in males (top) and females (bottom) from infected mothers, determined by RT-q-PCR. Each line connects two populations from same mouse. Statistics calculated by student’s paired t-test. (**C**) DotPlot showing differential variable chain gene transcription between *Xist* positive and *Xist* negative B cells from males from infected mothers. Average expression and dot size are as described above. (**D**) Fold change of *Xist* on bone marrow cells after *Xist* silencing, determined by RT-q-PCR. (**E**) Igα surface expression after *Xist* silencing *ex vivo*. Surface expression and cell population identity determined by flow cytometry. MFI normalized to average MFI of females from uninfected mothers for each experiment. Line connecting dots represent the same mouse. Statistics calculated by student’s paired t-test. (**F**) DiffScore of probes for UBL4A promoter probes from Illumina® Infinium HD Mouse Methylation Assay after ChIRP using probes for *Xist* normalized to females from uninfected mothers. Graph generated by Illumina® Genome Studio. (**G**) Fold change of *Ubl4a* on bone marrow B cell populations sorted by Igα high and low surface expression in males (top) and females (bottom) from infected mothers, determined by RT-q-PCR. Statistics calculated by student’s paired t-test. (**H**) Flow cytometric analysis of intracellularly stained phospo-STAT3 expression on whole bone marrow of pups from infected and control mothers. Experiment performed twice, representative of >3 litters. Statistics calculated by two-way ANOVA. (**I**) DotPlot showing differential expressed genes related to STAT3 pathway between *Xist* positive B cells from males and females from infected and control mothers. *p*-values on graph represent comparison between that group and males from infected mothers. Non-significant values (>0.05) not shown.

When comparing *Xist* positive and negative cells in males from infected mothers, *Xist* positive cells have increased transcription of various Ig heavy and light chain genes (Fig. 6C), suggesting differential V(D)J recombination. Moreover, some of these genes have been connected with autoimmune disease (*51*), of which aberrant *Xist* is known to play a pivotal role (*52-54*).

Overall, when comparing *Xist* positive cells to *Xist* negative cells in males from infected mothers, *Xist* positive cells downregulate multiple genes necessary for B cell function, such as *Cd79a, Cd79b, Ebf1, Rag1*, and *Cd74* (Fig. S6A). We then compared the *Xist* positive cells from males from control and infected dams. Males from infected mothers upregulate more genes, including those of functional importance for cells, such as *Il7r, Ikzf3*, and *Cd72* (Fig. S6B). This shows that not only are the *Xist* positive cell populations different between males from uninfected and infected mothers, but that the Xist positive cells from males from infected mothers are transcriptionally more diverse.

To examine the direct relationship between *Xist* expression in males from infected mother and Igα, *Xist* was silenced in whole bone marrow *ex vivo*, and developing B cell populations were analyzed. RT-qPCR was performed to confirm the downregulation of *Xist* expression (Fig. 6D). When *Xist* is silenced in males from infected mothers, Igα expression increases in both immature and T1 B cells (Fig. 6E). This is unique to males from uninfected mothers compared to females, suggesting normal *Xist* expression and regulation prevents this. Also, there is no change in Igα on males from uninfected mothers, most likely due to the rare amount of *Xist* positive cells, again suggesting atypical *Xist* expression in males causes decreased surface Igα.

From the ChIRP data, we determined that *Xist* is bound to the *Ubl4a* promoter, which is also highly methylated in males from infected mothers compared to other groups (Fig. 6F), suggesting lower transcript, as confirmed by RT-qPCR in Igα low, or *Xist* high B cells (Fig. 6G). Normally, UBL4A dephosphorylates STAT3 (*55*). STAT3 signaling is important for B cell maturation and function (*17, 56*). Due to the decrease in UBL4A, there in an accumulation of phospho-STAT3 in the bone marrow of males from infected mothers (Fig. 6H). Finally, in *Xist* positive cells, there is an increase in *Stat3* transcription and *Bcl2*, a STAT3-inducible gene, in males from infected mothers compared to males from uninfected mothers (Fig. 6I). This suggests an increase of activated, or phospho-STAT3, which cannot be dephosphorylated due to methylation of the *Ubl4a* promoter, causing an accumulation of phospho-STAT3 as seen in males from infected mothers at steady state and after immunization.

### Low Igα expression caused by high *Xist* expression leads to non-responsive B cells within the GC

To connect the developing B cell dysregulation to decreased humoral response, we employed adoptive cell transfers. Mature cell depleted bone marrow of males from control and infected dams were transferred to irradiated *Rag1*^*-/-*^ mice. After 5 weeks, the mice were given wildtype T cells and were immunized with Td a day later (Fig. 7A). Decreased GC B cells (Fig. 7B), decreased tetanus-specific B cells (Fig. 7C), and decreased ability to neutralize tetanus (Fig. 7D) were retained in recipients of bone marrow from males from infected mothers, confirming these defects in B cell development are sufficient to drive a dysfunctional humoral immune response.

**Figure 7:**
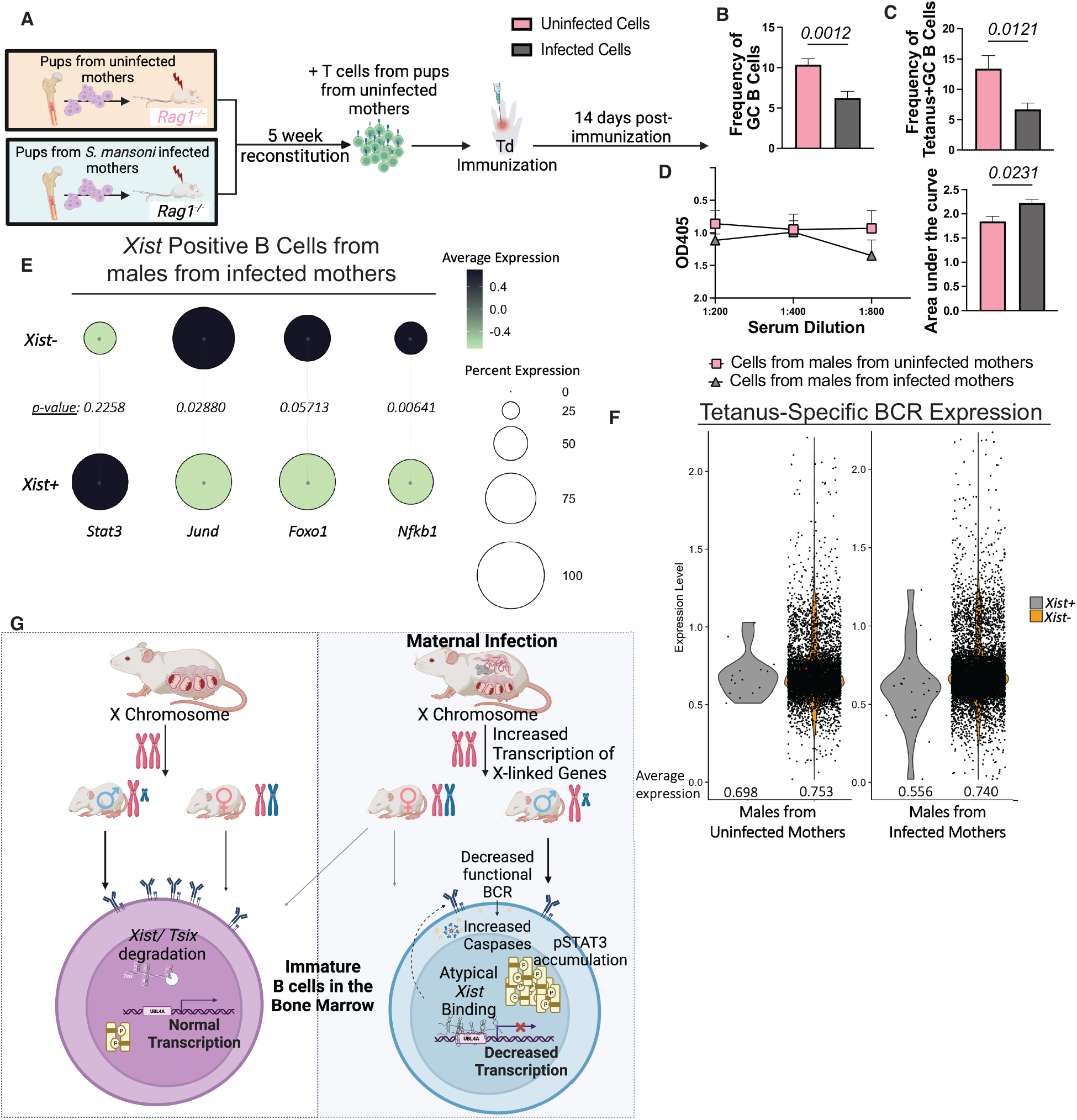
Dysfunctional Igα signaling caused by aberrant *Xist* expression in males from infected mothers lead to defective GC and recall response. (**A**) Schematic for adoptive transfer and immunization model. Flow cytometric frequencies of (**B**) GC B cells and (**C**) antigen-specific GC B cells from draining lymph node after adoptively transferring cells from control males and males from infected mothers to *Rag1*^*-/-*^ recipients before immunizing with Td and harvesting 14 days post immunization. Statistics calculated by student’s t-test. (**D**) Tetanus neutralizing assay from serum of *Rag1*^*-/-*^ recipients 14 days post immunization. Experiments repeated three independent times and represents >3 litters from infected mothers. (**E**) DotPlot showing genes related to B cell activation between *Xist* positive GC B cells from males from infected mothers. Average expression and dot size are as described above. (**F**) ViolinPlot showing expression level of tetanus-specific conjugated probe from single cell V(D)J sequencing of *Xist* positive and Xist negative GC B cells from the draining lymph nodes of male pups from infected mothers 14 days post Td vaccination. (**G)** Proposed schematic for model of maternal schistosomiasis induced dysfunction in B cell development.

We then looked at the effect of *Xist* in GC responses males from infected mothers. It has recently been published that *Xist* downregulation is important for the restraint of atypical B cells (*57*), but this work was done in females. When comparing the transcriptomes of *Xist* positive and negative cells within the GC of males (Fig. S7D, S7E), the atypical, or *Xist* positive cells, had a decrease in *Jund, Foxo1*, and *Nfkb1* (Fig. 7E), which transcription is all inducible by B cell activation and memory B cell formation (*58*), suggesting these B cells cannot effectively be activated due to defects in the BCR.

When comparing antigen specificity, *Xist* positive cells have, on average, a lower binding capability for tetanus compared to the *Xist* negative cells (Fig. 7F). Cells that bind labeled antigen to a lower extent have lower antigen affinity (*59*), showing that sustained *Xist* expression in the GC of males leads to an increase of low affinity B cells that would likely not mount a protective recall response.

## DISCUSSION

Although it is well established that maternal infections, specifically parasitic ones, can shape offspring immunity (*10-13, 60-63*), it is still unknown what factors influence offspring B cell development. Many current studies focus on TORCH (Toxoplasmosis, other (syphilis, varicella-zoster, parvovirus B19), Rubella, Cytomegalovirus, and Herpes) infections (*64*), or common congenital infections, with little effort on how non-congenital maternal infections affect offspring immunity. To understand how maternal infections alter immune development, it is necessary to not only characterize the defects in immunity that come with these infections, but also to understand the mechanisms underlying them. In this study, we uncover one mechanism behind decreased vaccine efficacy during maternal schistosome infection and link this to a sex-specific defect within bone marrow B cell development (Fig. 7G).

Starting with previously published clinical data (*14*), we found that male children from *S. mansoni* infected mothers have lower vaccine efficacy with no change in efficacy in females (Fig. 1F). Our established mouse model (*13*) not only mirrors this outcome but found this to be due to lower antigen specific B cells within the GC and a lower frequency of overall GC B cells in males from infected mothers only (Fig 1). It has been shown that high maternal antibody vaccine-induced titers can restrict offspring plasma cell generation and limit GC expansion to homologous immunization (*65*). Our model uniquely demonstrates that infection in dams, alters the GC response to heterologous immunization, suggesting another mechanism restricting GC function in pups from infected mothers. Additionally, while there is no functional antibody data in these children, we show that in mice, these antibodies generated have less tetanus neutralizing potential. Although there is an increased rate of outbreaks for vaccine preventable diseases in areas of Africa where helminths are endemic (*66*), this is attributed to more than just maternal infections, making it difficult to say to what extent maternal infections contribute to these outbreaks, but epidemiological data suggests maternal infections are playing a role.

We have connected this GC phenotype to developing cells within the bone marrow. Starting at the large pre-B cells stage, males from infected mothers have decreases in EBF1, *Id3*, and V_preB_ (Fig. 3A, 3B), which are essential for pre-BCR signaling and B cell development beyond this stage (*27, 28*). This then causes a dysregulation of pre-BCR signaling in males from infected mothers and a disruption in the developmental pathway. Pre-BCR signaling acts as a checkpoint within B cell development (*67*), so when this fails, as seen in an increased number of pre-B cells in males born to infected mothers, there is a decrease in the immature B cell and subsequent T1 B cell pools (Fig. 3D, 3E).

ScRNASeq revealed that males from infected mothers have an overall transcriptional decrease in machinery needed for receptor editing. Receptor editing is a secondary method of V(D)J recombination specific to the light chain to optimize a BCR after failed tolerance (*34, 35*).

Roughly 25% of B cells undergo receptor editing (*68*), which matches the loss in naïve B cells seen in the periphery compared to T1 B cells in the bone marrow of control pups (Fig. 2A). Yet, because of the absence of the ability to undergo receptor editing, we see that males from infected mothers often fall below the best fit line (Fig. 2A), suggesting a loss of greater than 25% of B cells in the periphery, resulting in a lower pool of peripheral B cells available during infection or immunization.

In conjunction with lower pre-BCR signaling potential, we also observed a decrease in Igα on immature and T1 B cells in males from infected mothers (Fig. 3G, 3H). Igα is one the signaling components of the BCR and is necessary for BCR signaling during selection and the GC reaction (*34, 38*). In humans, a deficiency in Igα results in a complete block in B cell development (*38*). It must be noted that these cells do not lose all Igα surface expression, but a proportion of these B cells have a decrease in the amount of Igα on the surface, resulting in a population of B cells with reduced capacity to signal through the BCR. Although little is known about the signaling that occurs when Igα is downregulated and not complexly abolished, it is hypothesized that decreases in BCR signaling strength during the immature and the T1 stage determine survival (*69*). We found, through adoptive transfers, that this is cell intrinsic, rather than being caused by the microenvironment, concretely connecting this developmental defect to the decrease in peripheral B cells (Fig. 3J-N). There is known crosstalk between peripheral B cells and the bone marrow to regulate the rate of lymphopoiesis (*70*), but this was demonstrated in the complete absence of peripheral B cells. It is unknown what triggers this response and if there is a threshold of B cell loss that is needed. Our data suggests this to be true because there is not increased lymphopoiesis in our model, indicating that either this feedback mechanism is also defective or the threshold to increase lymphopoiesis has not been reached.

Using single cell V(D)J sequencing of bone marrow B cells, we found males from infected mothers have an expansion of unique clones that are not shared with their littermate female counterparts or males from control mothers (Fig. 4A, 4C). This can likely be attributed to variations in λ gene usage. Males from infected mothers readily pair *Iglv2* and *Iglj3* in addition to *Iglv1* and *Iglj2* (Fig. 4D). Both pairings are absent in other groups, indicating males from infected mothers can make unique light chain gene pairings. While we cannot say if these pairings are functional or not, it can be assumed that if they were highly functional, they would also be enriched in other groups. These likely also contribute to the increased expression of genes related to the Caspase pathways (Fig. 4E). Together, these pairing defects contribute to the decrease in peripheral B cells.

Thus far, much of this data presented has the potential to be infection specific, or only seen within maternal schistosome infection. But we are the first to characterize relatively high *Xist* expression in males due to methylation differences on the X chromosome (Fig. 5). This circumstance is relevant to multiple fields and studies, such as in models of lupus (*52, 54*), certain cancers (*71, 72*), and other autoimmune diseases (*73*). As shown in this study, aberrant *Xist* expression in males causes a decrease in surface Igα expression due to an accumulation of pSTAT3 (Fig. 6), a feature seen in models of lupus (*74-76*). While it is unknown why males who have systemic lupus erythematosus have aberrant *Xist* expression (*52, 54*), this study suggests maternal factors can contribute to this phenotype and may cause disease later in life. Future work will further explore this relationship.

Our previous work established that peripheral follicular dendritic, T follicular helper, and GC responses were all diminished in offspring born to infected mothers. These new findings link defects in B cell development cell-intrinsically to defects in the GC reaction. By adoptively transferring bone marrow to irradiated recipients with control T cells, we show that defective B cell development can cause a decrease in antigen specific B cells despite normal T and stromal cell populations (Fig. 7B-D). B cells with high *Xist* expression within the GC also have a decrease in transcriptional activation indicators (Fig. 7E) and a decrease in antigen affinity (Fig. 7F). Recent publications have demonstrated the importance of *Xist* within GC B cells, stating that *Xist* is upregulated when activated (*45*) to restrain atypical memory B cell formation (*57*), which have been associated with disease (*77*). These studies have only used female subjects. It is unknown if this also occurs in males, and is speculated not to be, since atypical memory B cells do not accumulate in healthy males (*78*). Our model demonstrates that aberrant increased *Xist* expression in males is sustained from B cell development through the GC reaction, causing defects in BCR signaling following maternal infection.

Other maternal infection models studying offspring neurological effects and perturbations within offspring gut microbiota have shown IL-6 as the main culprit for these effects (*79-81*), including lower vaccine efficacy in mice (*82*), this is not true during maternal schistosomiasis. While excessive maternal IL-6 can lead to increased IL-6 and IL-17A in offspring, this is not seen in our model (Fig S8B, S8C), suggesting there is more than just IL-6 exposure involved.

Additionally, epidemiological data has implicated maternal genomic imprinting in altered offspring immunity (*83*), and maternal X chromosome and *Xist* expression has been demonstrated to be able to be imprinted in males (*84-86*). In our model, schistosome infected mothers do have increased *Xist* expression, as shown by flow-fish (Fig. S8C). This is not surprising because *Xist* can be upregulated by active infection or inflammation (*87*). We postulate that this increase in *Xist* expression due to infection may be imprinted on the offspring’s maternal chromosome, inappropriately increasing *Xist* expression in offspring, especially in males.

This study was limited by the chosen challenge model. Immunization models allow for antibody titer and neutralizing capacity as outcome measures and a clear phenotype to characterize. Because of the combined Td immunization used, the only challenge model is a lethal tetanus toxin challenge with limited ability to parse nuances in immunological phenotype.

Future studies will utilize our mouse model and human studies to further look at maternal schistosomiasis and offspring immunity. Using the mouse model, varying vaccination techniques and adjuvants to increase vaccine efficacy after maternal infection will be tested. Additionally, further work to understand stepwise B cell selection and tolerance in the bone marrow and the nature of the unique clones that occur in males from infected mothers is needed. Most of this work will need to be validated in humans during maternal schistosome infection using the outcome measures identified in this work in peripheral blood B cells.

Overall, these data are the first to show a male-specific role of *Xist* expression and dysregulation and provide new insight on potential targets for therapeutics to improve vaccine efficacy and lower childhood morbidity.

## Supplementary Materials

Materials and Methods

Figs. S1 to S8

References (88-100)

## Acknowledgments

B. glabrata snails provided by the NIAID Schistosomiasis Resource Center of the Biomedical Research Institute (Rockville, MD) through NIH-NIAID Contract HHSN272201700014I for distribution through BEI Resources. We thank Dr. Dan Colley for sharing the unpublished data from the Kenyan maternal schistosomiasis trial, and Dr. Montserrat C. Anguera for providing all the reagents and protocols need for RNA FISH.

## Author contributions

Conceptualization: LCG, JMO, KCF

Data curation: LCG, JMO, BNO, KCF

Formal analysis: LCG, KCF

Funding acquisition: KCF

Investigation: LCG, JMO, KCF

Methodology: LCG, JMO, KCF

Project administration: KCF

Resources: KCF

Supervision: KCF

Validation: KCF

Writing – original draft: LCG, KCF

Writing – review & editing: LCG, JMO, BNO, KCF

## Data and materials availability

All requests for resources and reagents should be directed to and will be fulfilled by the Lead Contact Keke Fairfax. Single-cell RNA-seq data have been deposited at GEO and are publicly available as of the date of publication. Accession numbers will be available after paper acceptance for publication. Scripts for scRNAseq analysis are deposited on GitHub and will be made public after paper acceptance for publication. Microscopy data reported in this paper will be shared by the lead contact upon request.

