## Supplementary material for "Maternal infection causes dysfunctional BCR signaling in male offspring due to aberrant Xist expression": methods and supplementary figures

### MATERIALS AND METHODS

#### *Study Design*

This study aimed to understand mechanisms that underlie decreased vaccine efficacy during maternal schistosomiasis, specifically focusing on the B cell response. Using our previously established maternal schistosomiasis murine model (*13*), we generated pups from *Schistosoma mansoni* infected and sham infected mothers. Sample size for each experiment was limited by liter size. The number of independent experiments is indicated in the figure legend. Pups were excluded if the dam stopped breastfeeding the pups before 14 days of age, determined by weanling weight. Outliers were excluded by the ROUT method with Q = 0.1% to ensure only definitive outliers were removed.

To determine developmental B cell defects in pups from *S. mansoni* infected mothers that could lead to a diminished vaccine response, flow cytometry and single cell RNA sequencing were coupled in the bone marrow and peripheral lymph nodes. V(D)J sequencing was performed to establish if maternal infection also altered the homeostatic and germinal center B cell repertoires. Due to the sex-specific nature of the results, and an increase of *Xist* expression in males from infected mothers, we performed a pulldown using biotinylated probes for *Xist* and extracted the DNA for a methylation array. Silencing of *Xist* in whole bone marrow *in vitro* determined the relationship between dysfunctional *Xist* expression and the BCR development. Finally, adoptive transfers were conducted to test the origin, longevity, and persistence of these defects. Experiments were not blinded.

#### *Mice*

4get homozygous ($Il4^{tm1Lk}$)(strain# 004190) mice and $Rag1^{-/-}$ (C.129S7(B6)-*Rag1$^{tm1Mom}$*/J)(strain# 003145) mice were obtained from The Jackson Laboratory and bred at the University of Utah animal facilities. Experiments were performed in strict accordance with the NIH guide for the Care and Use of Laboratory animals and institutional guidelines for animal care at the University of Utah under approved protocols (18–09001 and 21-07009). At six weeks of age, 4get homozygous females were infected with infected with ~35 *Schistosoma mansoni* cercaria, as previously described (*13*), or sham infected. Six weeks post *S. mansoni* or sham infection, females were paired to KN2 homozygous ($Il4^{tm1(CD2)Mmrs}$) (*16*)) males and remained paired until they reached humane endpoints. To generate 4getKN2 and KN2 homozygous littermates, 4getKN2 females were infected and mated to KN2 homozygous males as described above. Pups from mixed genotype litters were genotyped by flow cytometry before use.

*In vivo treatments*

Pups were weaned and genotyped (as necessary) at 28 days old. Steady state experiments were performed between 28-35 days of age. For immunization experiments, pups were immunized with 1/10$^{th}$ the human dose of Tetanus/Diphtheria commercial vaccine (Grifols Therapeutics Inc (TDVAX) #13533013101) subcutaneous in the rear footpad. Mice were sacrificed at 14-days post-primary immunization or administered a secondary immunization at over 60 days post-primary immunization and sacrificed 3 days post-secondary immunization. For all experiments, littermates were used, and groups were split by sex and infection status of the mother.

*Cell isolation for single cell suspension*

For bone marrow harvesting, femurs were collected, and the tip connected to the knee was clipped on a bias to expose the bone marrow. Bones were placed cut side down into a 200µL microcentrifuge tube with a hole in the bottom and ~50µL of Dulbecco's Modified Eagle Medium (DMEM)(Corning #10-013-CV). This tube was then placed into a 1.5mL microcentrifuge tube and then pulsed in a microcentrifuge at ≥ 10,000 RPM for 15 seconds at 4°C. The 200µL tube with the flushed bones were then discarded. Erythrocytes were lysed using 1X lysis buffer (10X BD Pharm Lyse™, BD Biosciences #555899)(diluted to 1X) for 1 minute. The lysis was quenched using 1% fetal bovine serum (FBS)(Gibco #26140-079) in DMEM and centrifuged at 200xg for 5 minutes at 4° C. The supernatant was discarded and washed with 1X phosphate-buffered saline (PBS) before cell surface staining.

Popliteal lymph nodes (PLNs) and blood were collected and processed as previously described (*88*). In brief, lymph nodes were collected, smashed through a 100µm cell strainer, and washed with 15mL of DMEM.

*Flow cytometry and antibodies*

To stain cells for flow cytometry, cells were harvested as described above and resuspended in the appropriate volume of FACS buffer (2%FBS, 5mM EDTA in 1X PBS) as previously described (*13, 88*). The following antibodies from Invitrogen were used for cell surface staining: CD34 - Biotinylated (RAM34), CXCR5 Biotinylated (SPRCL5), CD150 Super Bright 600 (mShad150), CD150 PerCP-eFluor710 (MShad150), CD24 Super Bright 600 (M1/69), CD19 Super Bright 780 (1D3), CD95 AF488 (15A7), CD11c FITC (N41B), CD3 FITC (17A2), Ter-119 FITC (TER119), B220 PerCP-Cy5.5 (RA36B2), Sca-1 APC (D7), CD48 AF700 (HM48-1), IgM PE-Cy7 (II/41). The following antibodies from Biolegend were used for cell surface staining:

Zombie Red™ Dye (for viability), Streptavidin APC/Fire™ 750, CD135 Brilliant Violet 421™ (A2F10), CD135 Brilliant Violet 421™ (A2F10), CD127 BV711 (A7r34), GR-1 FITC (RB6-8C5), B220 FITC (RA36B2), CD11b FITC (M1/70), CD79a APC (F11-172), CD179a PE (R3), CD16/32 PE-Cy7 (93). The following antibodies from BD Biosciences were used for cell surface staining: IgD BV510 (11-26c.2a), CD117 BV650 (2B8), CD43 BV650 (S7), GL7 AF647 (GL7), GL7 PE (GL7), CD117 PE (2B8). For transcription factor staining, cells were fixed and permeabilized with the FoxP3 transcription factor staining kit (Invitrogen # 00-5523-00) per manufacture's instruction, and subsequently stained with combinations the following transcription factor-specific antibodies from BD Biosciences (EBF1 PE (T26-818) and Caspase-3 BV650 (C92-605)) and Invitrogen (ID2 APC (ILCID2), Ki67 APC (SolA15), and Phospho-Stat3(Tyr705) eFluor450 (LUVNKLA)).

Bone marrow B cells gating was done after first gating out Dump$^{+}$ cells (Viability$^{+}$CD3$^{+}$CD4$^{+}$CD8$^{+}$CD11b$^{+}$CD11c$^{+}$Gr-1$^{+}$Ter119$^{+}$). Pre–pro-B cells (B220$^{+}$CD19$^{-}$), pro-B cells (B220$^{+}$CD19$^{+}$IgM$^{-}$IgD$^{-}$CD43$^{+}$), large pre-B cells (B220$^{+}$CD19$^{+}$IgM$^{-}$IgD$^{-}$CD43$^{+}$FSC$^{hi}$), small pre-B cells (B220$^{+}$CD19$^{+}$IgM$^{-}$IgD$^{-}$CD43$^{+}$FSC$^{lo}$), immature B cells (B220$^{+}$CD19$^{+}$IgM$^{+}$IgD$^{-}$), and transitional B cells (B220$^{+}$CD19$^{+}$IgM$^{-}$IgD$^{+}$) gating was adopted previously published studies (*21-23*). PLN germinal center (GC) B cells are described as Viability$^{-}$CD19$^{+}$GL-7$^{+}$FAS$^{+}$. All flow cytometry experiments were run on an Attune NxT (Invitrogen), and data were analyzed with FlowJo.

*Tetanus neutralization assays*

Modified tetanus neutralization assays(*89*) were performed by first coating an Immulon 4 HBX 96 well plate (Fisher Scientific #14-245-153) with 10μg/mL of trisialoganglioside-$GT_{1b}$ from bovine brain (Sigma-Aldrich # G3767-1MG) in ethanol overnight. The next day, the plate was blocked with 1% BSA for two hours at room temperature. Meanwhile, serum from Td immunized pups were incubated with tetanus toxin for two hours at 37°C. After flicking off the blocking solution, the serum incubated with tetanus was added to the plate and let to sit at room temperature for two hours. After washing, anti-tetanus toxoid rabbit serum (VWR # BOSSBS-11772R) was added to the plate for two hours at room temperature. After washing, anti-rabbit IgG HRP (Abcam # ab6721) was added to the plate and left to incubate for two hours at 37°C. Finally, SuperAqua Blue ELISA substrate (Thermo Fisher Scientific #00-4203-56) was added to each well before reading plate at 405nm with spectrophotometer.

*Adoptive cell transfers*

Steady state experiments were performed by harvesting whole bone marrow from male 4getKN2 pups from infected and control mothers as described above. Cell suspensions were pooled before counting using a hemocytometer. Three million cells were then injected intravenously into 4–6-week-old Rag1$^{-/-}$ recipients. Recipients were bled weekly to check lymphocyte reconstitution. At 5 weeks post-transfer, recipients were euthanized, and cells were collected for flow cytometric analysis.

Immunization experiments began by partially irradiating Rag1$^{-/-}$ with 300 grays. The following day, donor cells were collected from bone marrow of males from infected and control mothers.

Mature cells ($CD3^+CD4^+CD8a^+B220^+CD11b^+CD11c^+Gr\text{-}1^+Ter119^+$) were depleted by using the EasySep™ FITC Positive Selection Kit II (Stem Cell #17682) per manufacturer's instruction. Non-labelled cells were then counted using a hemocytometer and $5x10^5$ cells were transferred intravenously into each $Rag1^{-/-}$ male recipient mouse. Recipients were allowed 5 weeks to reconstitute before transferring $5x10^5$ T cells from wildtype 4getKN2 females. The following day, mice we immunized in the footpad with $1/10^{th}$ the human dose of the commercial Tetanus/Diphtheria vaccine listed above. Fourteen days after immunization, mice were euthanized, and cells were harvested for analysis.

*Tetanus toxin fluorophore conjugation*

Tetanus toxin (List Labs #190B) was conjugated to using the Lightning Link ® FITC conjugation kit (Abcam #ab102884) per manufacturer instruction. Ovalbumin conjugated to Biotin (Abcam #ab201795) was used in conjunction with the conjugated Tetanus toxin to control for nonspecific B cell binding. The decoy was incubated with streptavidin-BV786 (BD Biosciences #563858) for 10 minutes at RT before mixing with the conjugated Tetanus toxin antibody. Cells were stained with 0.01µL of conjugated Tetanus toxin for 20 minutes 4° C and washed twice with 1X PBS before surface antibody staining.

*scRNASeq experiments*

Single cell suspensions were isolated from tissue as described above then sorted using either a FACSAria or a Sony MA800 or MA900 cell sorter. Cells were processed as previously described (*13*). In brief, sorted cells were pelleted by centrifugation and resuspended in 1% BSA (bovine serum albumin) in 1X PBS and counted by hemocytometer and resuspended to a concentration of 1,200 cells/μL to run on the 10x platform. Paired-end RNASeq (125 cycles) was performed via an Agilent HiSeq next-generation sequencer.

*scRNASeq analysis*

Raw data from scRNASeq experiments in this manuscript can be found in the NCBI's Gene Expression Omnibus database. Sequencing reads were processed by using a 10x Genomics CellRanger pipeline and further analyzed using R Studio. Prior to analysis, low quality cells (greater than 15% mitochondrial gene representation and/or fewer than 200 and/or more than 6000 genes per cell) were filtered out before clustering and differential expression testing (DEGs) by using the Seurat package (*90-92*). The biological identities of cell clusters were annotated by referencing a published scRNASeq data set(*93*) and by surveying known transcriptional cell markers in the scRNASeq data set.

*V(D)J scRNASeq and analysis*

Cell isolation, cell sorting, single cells clustering, and analysis were performed as described above. Data sets were analyzed using both scRepertoire (*94*) and VDJView (*95*).

*Xist RNA fluorescent in situ hybridization*

CLPs (viability$^-$CD3$^-$CD4$^-$CD8$^-$B220$^-$CD11b$^-$CD11c$^-$Gr-1$^-$Ter119$^-$cKit$^+$Sca-1$^+$Flt-3$^+$IL-7R$\alpha^+$(*96, 97*) were isolated and sorted using the strategies outline above. After sorting, cells were cytospun at 400xg for 5 minutes onto microscope slides (VWR #48311-703). Slides were prepared and stained using established protocols (*45*). Two Cy3-labeled 20-nucleotide oligo probes were designed to recognize regions within exon 1 (a gift from Montserrat Anguera). Slides were imaged at 63x using a Leica SP8 DIVE multiphoton microscope.

*Xist chromatin isolation by RNA purification (ChIRP)*

Whole bone marrow was isolated from femurs as described above. Cells from 3-4 mice were pooled before further processing. The RNA pulldown was done using a previously established protocol (*98, 99*) with the following adaptations; 1) cells were crosslinked with 1% formaldehyde for 20 minutes; 2) after hybridization and washing, DNA was isolated using PhOH:Chloroform:Isoamyl (Invitrogen #15593-031).

*Illumina® Infinium HD Mouse Methylation Assay*

Bisulfite conversion of DNA was performed using the EZ DNA Methylation-Gold kit (Zymo, catalog #D5006) as per manufacture's instruction. Infinium HD methylation microbead array (#20041558) was performed by the DNA sequencing core at the University of Utah following manufacturer's protocols. Data was processed and analyzed using the Illumina GenomeStudio Software.

*Enzyme-linked immunosorbent assay (ELISA)*

ELISA data used was published in (*13*), but split by sex of the pups and reanalyzed. Avidity ELISA were performed by following the above ELISA protocol with an additional 15-minute agitated incubation with 0.5M ammonium thiocyanate for treated wells or 15-minute incubation with 1X PBS for control wells before secondary staining as described in (*100*).

*RNA isolation and Quantitative polymerase chain reaction*

RNA isolation and q-RT-PCR were performed as previously described (*88*). In short, 30,000-50,000 cells were sorted into 500µL of TRIzol™ LS (Invitrogen # 10296010) and stored at -80° C until RNA extraction as previously described (Fairfax, 2013 #271). RNA was used for cDNA synthesis using Superscript IV VILO (Invitrogen # 11756050) for qPCR analysis. qPCR was performed using TaqMan™ Gene Expression Master Mix (Applied Biosystems #4369016) as per manufacturer's instruction with Xist probe (Mm01232884_m1)(Thermo Scientific #4331182) and beta actin probe (Mm02619580_g1)(Thermo Scientific #4331182) as the housekeeping gene. q-RT-PCR was run on an Applied Biosystems QuantStudio™ 3 Real-Time PCR System.

*Serum Cytokine Measurements*

Serum collected from pups 28-35 days after birth and 14 days post-immunization were collected upon sacrifice and stored at -80° C until use. Each grouping represents 2-3 different litters/experiments. The Cytokine & Chemokine Convenience 36-Plex Mouse ProcartaPlex™ Panel 1A (Thermo Fischer Scientific # EPX360-26092-901) was used per manufacturer instruction using a Luminex Magpix system as previously described (Cortes-Selva et al., 2021a). MFI was analyzed using the ProcartaPlex Analyst 1.0 Software.

*Flow-FISH*

Peripheral blood mononuclear cells (PBMCs) were isolated from whole blood of control and *S. mansoni* chronically infected 4get females. Whole blood was lysed using three times using 1X diluted lysis buffer (BD Biosciences #349202). Xist probes used were the same as used for

RNA FISH. Surface staining, fixation, permeabilization, and Xist staining was performed as described by (Arrigucci et al., 2017).

*Computer software versions*

R_4.2.1, dbplyr_2.2.1, devtools_2.4.4, dplyr_1.0.1.0, edgeR_3.39.6, ggplot2_3.3.6, ggrepel_0.9.1, HDF5Array_1.25.2, hdf5r_1.3.5, RColorBrewer_1.1-3, rhdf5_2.41.1, scRepertoire_1.7.2, sesame_1.15.8, Seurat_ 4.1.1, shiny_1.7.2, and wesanderson_0.3.6 were used.

*Statistical Analysis*

For scRNASeq experiments, *p*-values and adjusted *p*-values were calculated while doing differential gene expression analysis, which uses a non-parametric Wilcoxon rank sum test. All other statistical analyses were done using GraphPad Prism v8.0. For multiple comparisons, data was analyzed by analysis of variance (ANOVA) with Turkey's multiple comparisons test. *p*-values $\leq 0.05$ were considered statistically significant and is denoted in figures.

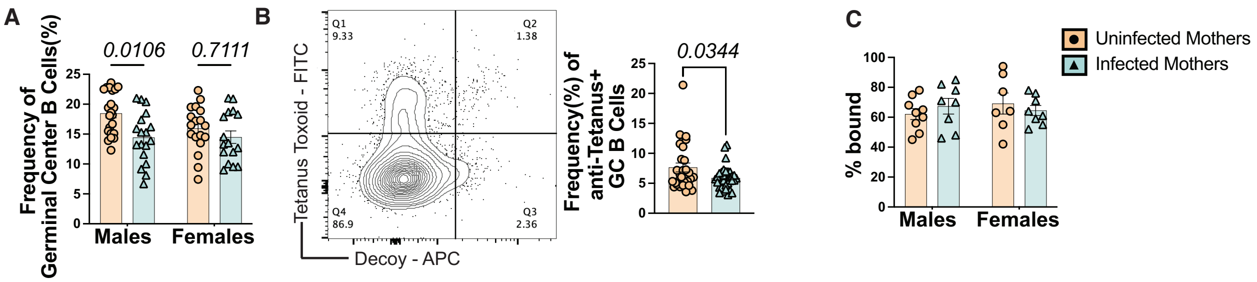


**Figure S1: Pups from *S. mansoni* infected mothers have decreased antigen-specific B cells from germinal center reaction.** (**A**) Flow cytometric frequencies of GC B cells ($CD19^{+}CD95^{+}GL7^{+}$) from the draining lymph nodes of pups from control and infected mothers 14 days post Td vaccination. Statistics calculated by two-way ANOVA. (**B**) Concatenated flow cytometry plot showing efficiency of conjugated tetanus antibody compared to a conjugated non-specific antigen from the draining lymph node 14 days post Td vaccination (left) and quantification of Q1 (right) that represents tetanus-specific GC B cells. Statistics calculated by student's t-test. (**C**) Antibody affinity ELISA done using serum from pups from control and infected mothers 14 days post Td vaccination. Statistics calculated by two-way ANOVA.

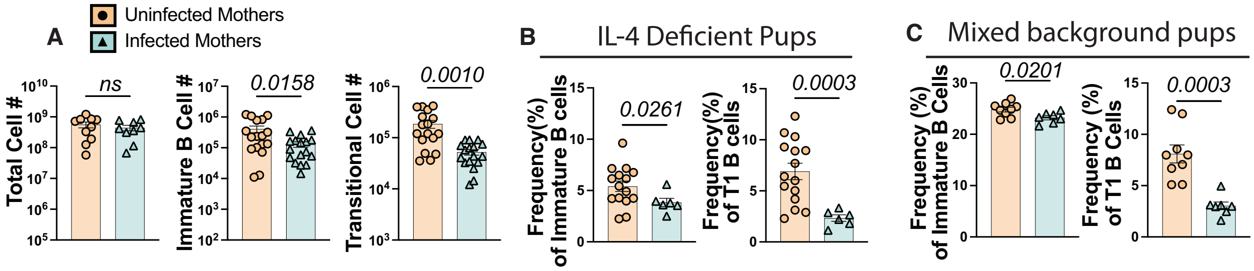


**Figure S2: B cell development is altered during maternal schistosomiasis independent of IL-4 and mouse background.** (**A**) Total cell numbers from bone marrow (left), immature B cells (middle), and T1 B cells (right) from the bone marrow of pups from control and *S. mansoni* infected mothers. Experiments performed >3 times, representative of >6 litters. (**B**) Flow cytometric frequencies of immature (left) and T1 (right) B cells in IL-4 deficient KN2 homozygous pups from *S. mansoni* infected mothers. Experiments performed twice, representative of >3 litters. (**C**) Flow cytometric frequencies of immature (left) and T1 (right) B cells from the bone marrow of pups born to control and *S. mansoni* infected CD45.1 mothers bred to Balb/cJ fathers. Experiments performed twice, representative of >4 litters. Statistics determined by student's t-test.

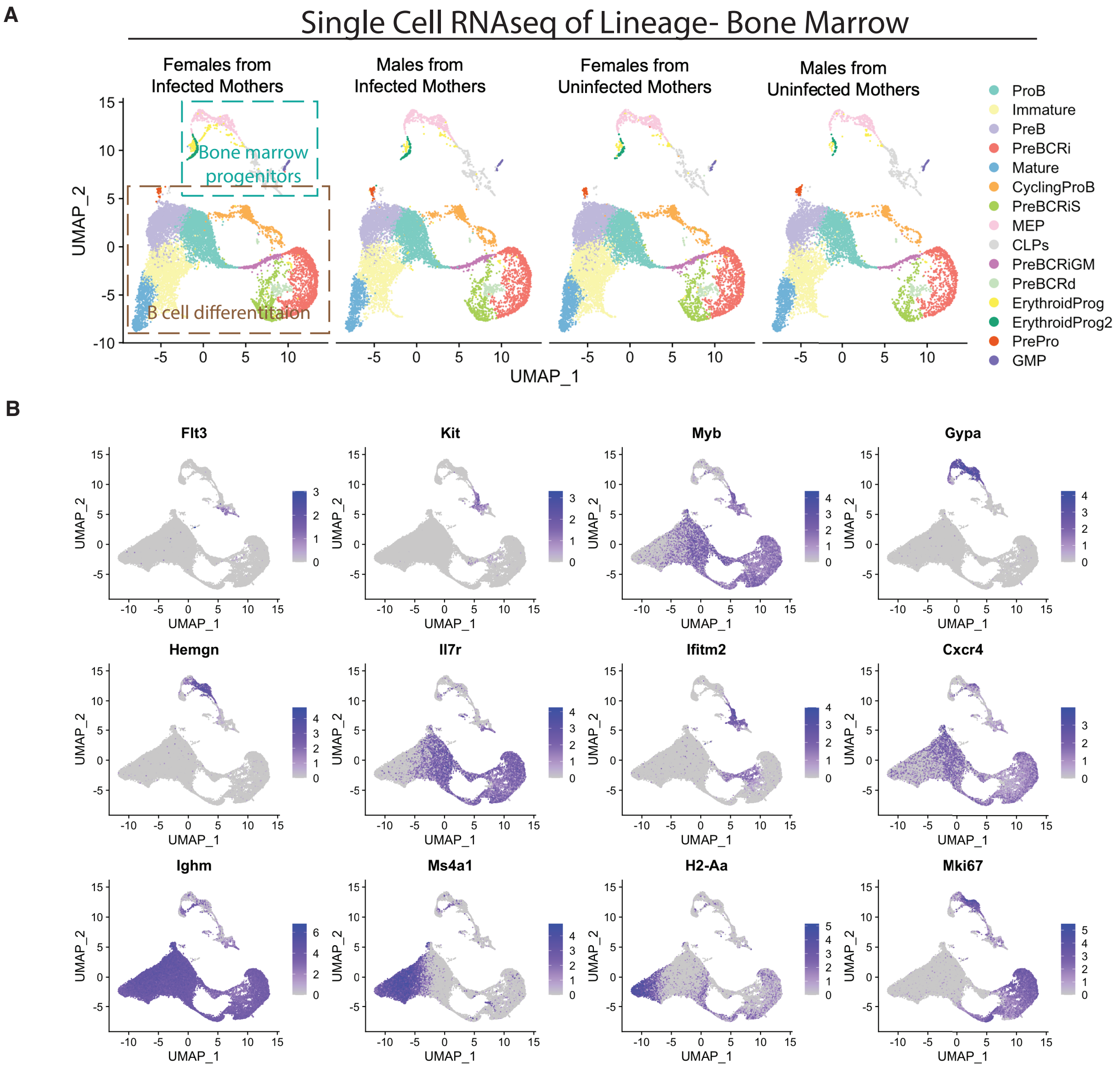


**Figure S3: Single cell RNA sequencing of myeloid depleted bone marrow from pups from *S. mansoni* infected mothers.** (**A**) UMAP showing labeled clusters from single cell RNA sequencing of bone marrow from pups from control mothers and mothers infected with *S. mansoni*. Each group represents a pool of >4 mice. (**B**) Transcriptional markers shown as FeaturePlots used to identify different clusters.

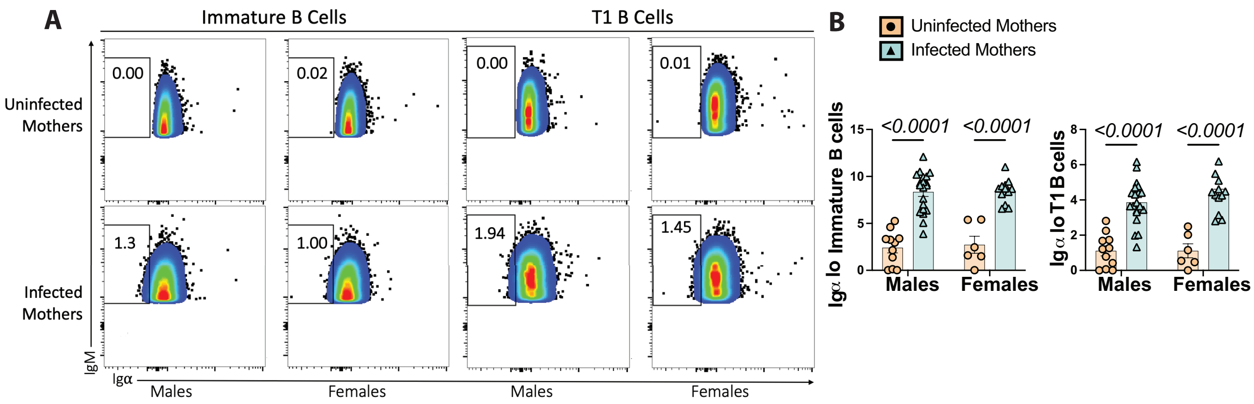


**Figure S4: Igα surface expression on immature and T1 B cells from pups from *S. mansoni* infected mothers is decreased.** (**A**) Concatenated flow cytometry plots of Igα staining on immature and T1 B cells. (**B**) Frequencies of gated Igα low cells in pups from control and infected mothers on immature (left) and T1 (right) B cells. Statistics calculated by two-way ANOVA.

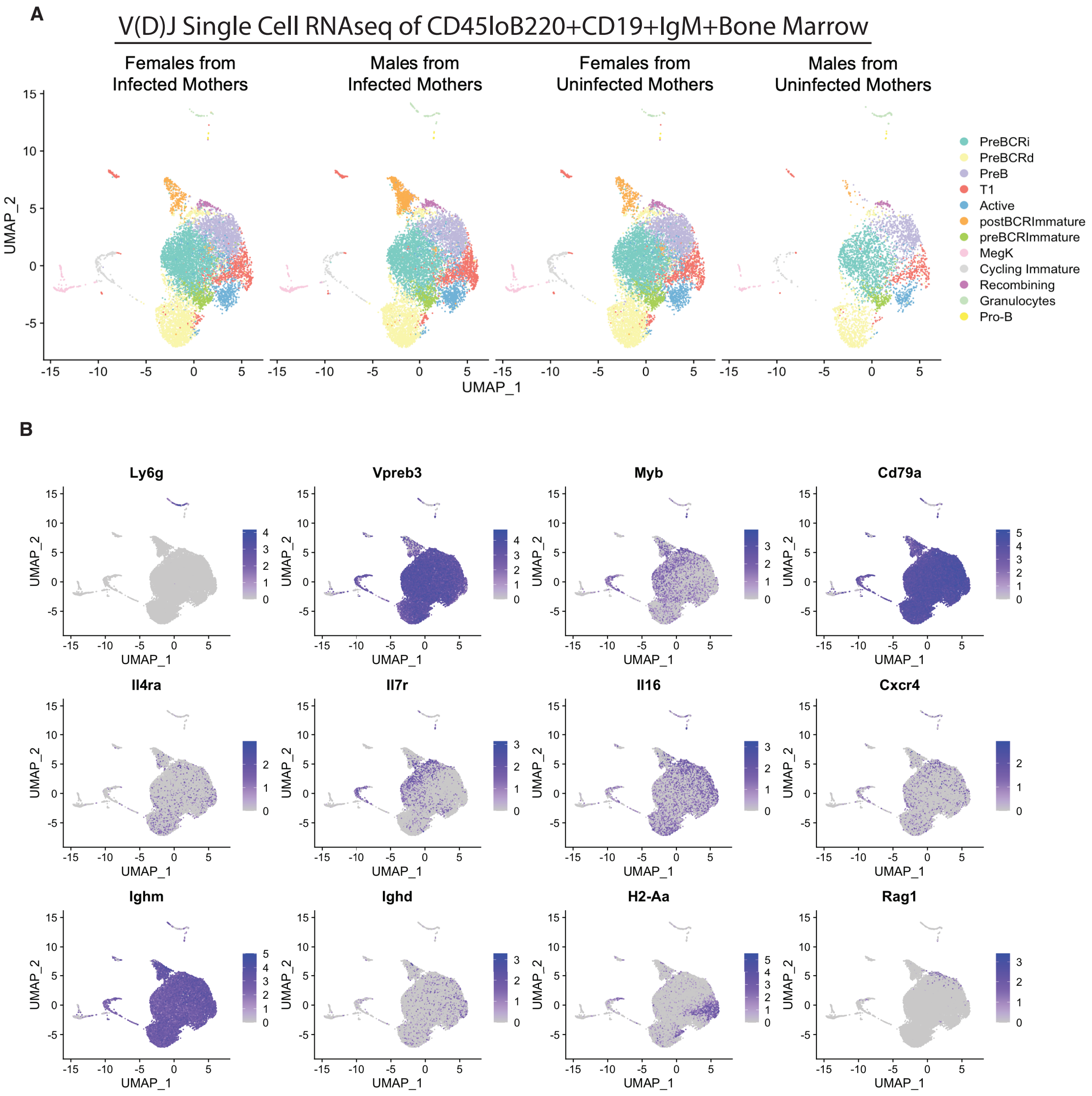


**Figure S5: Single cell V(D)J sequencing of immature and T1 B bone marrow from pups from *S. mansoni* infected mothers.** (**A**) UMAP showing labeled clusters from single cell RNA sequencing of bone marrow from pups from control mothers and mothers infected with *S. mansoni*. Each group represents a pool of >4 mice. (**B**) Transcriptional markers shown as FeaturePlots used to identify different clusters.

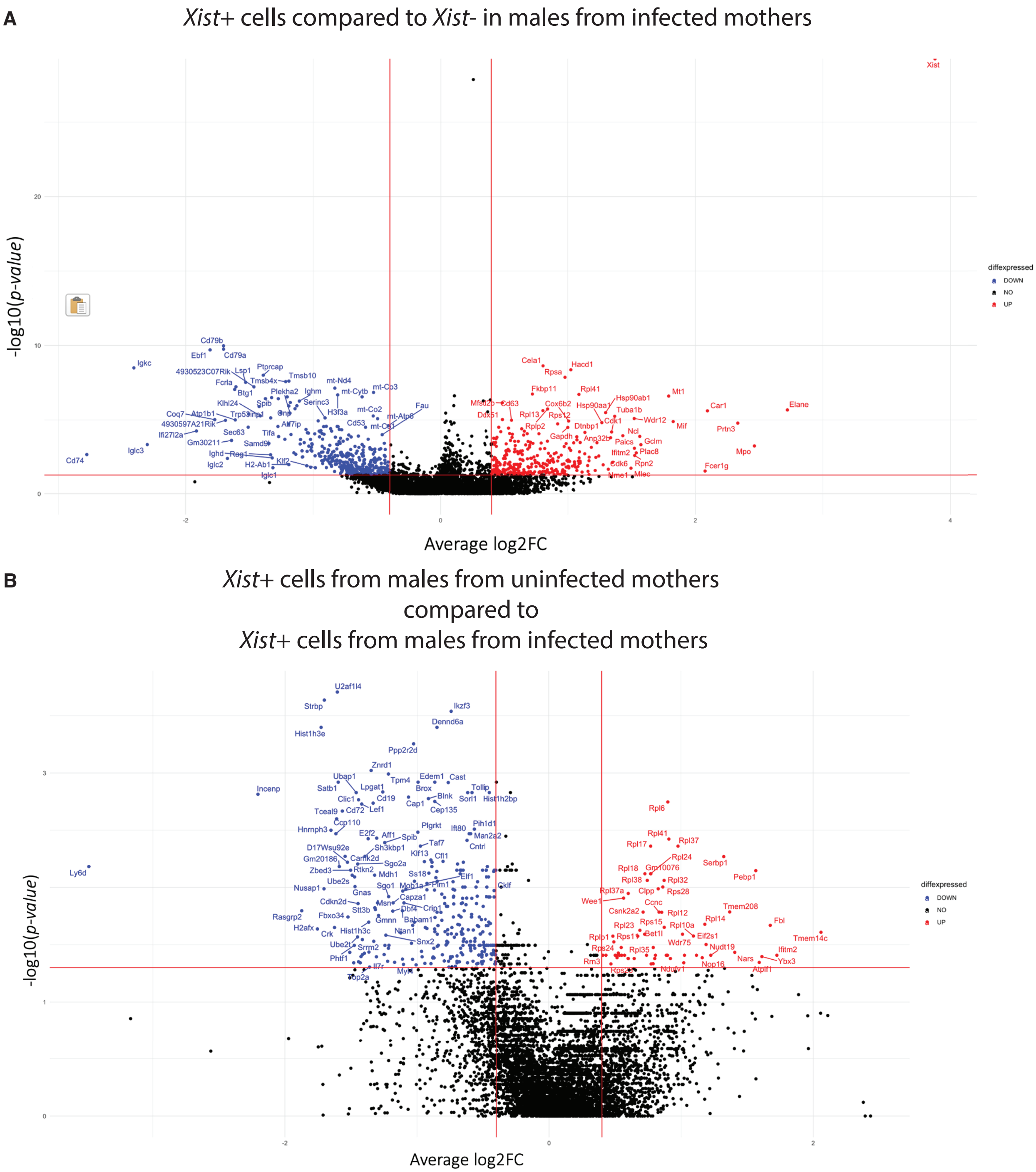


**Figure S6: *Xist* positive B cells in males from *S. mansoni* infected mothers are transcriptionally distinct from *Xist* negative B cells and *Xist* positive B cells from males from control mothers.** Volcano plots showing differentially expressed genes comparing (**A**) *Xist* positive B cells to *Xist* negative B cells in males from infected mothers and (**B**) *Xist* positive B cells between males from uninfected mothers and males from infected mothers from single cell RNA sequencing shown in Figure S3.

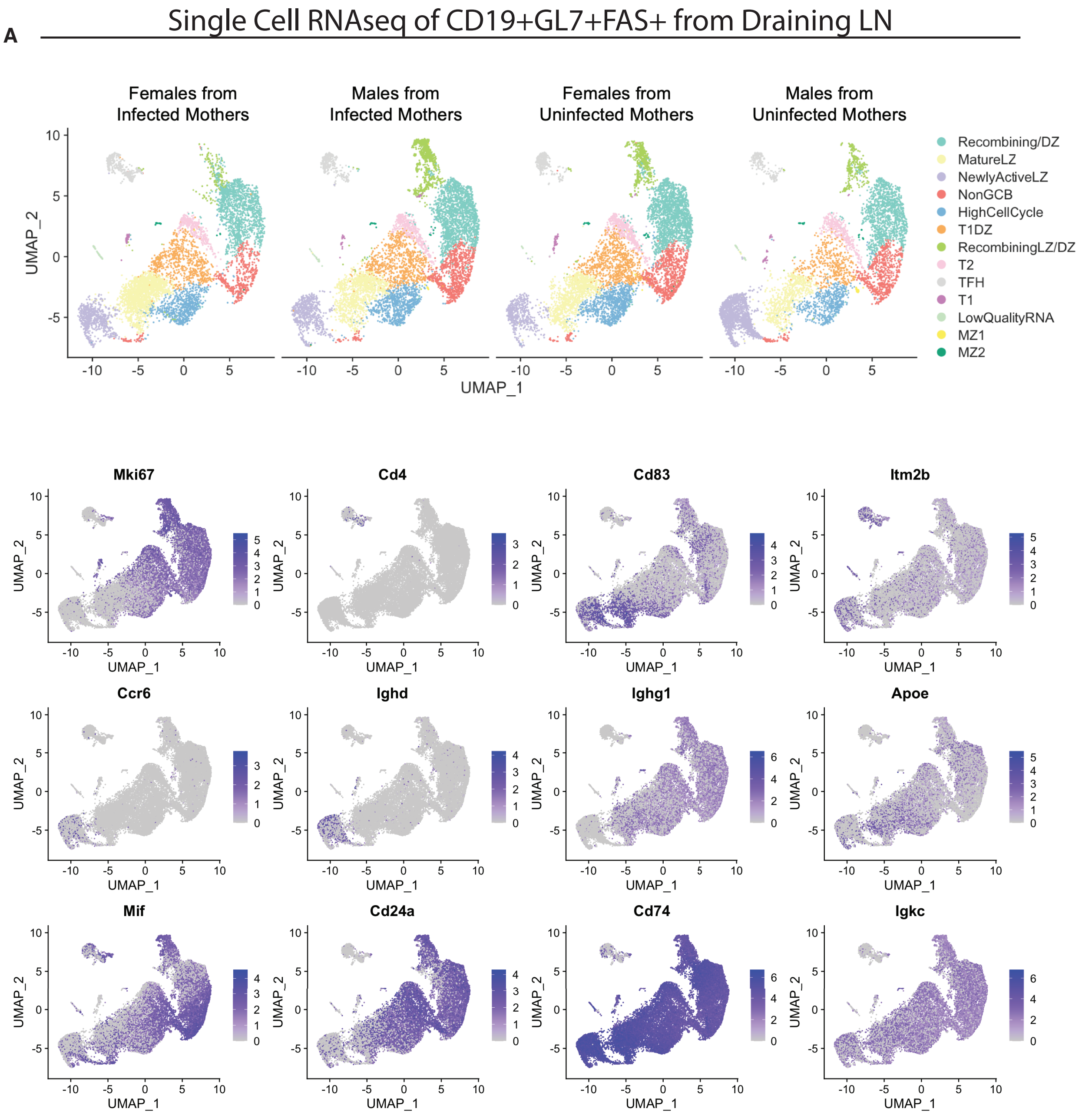


**Figure S7: Single cell V(D)J sequencing of draining lymph node after vaccination. (A)** UMAPs generated after sorting $CD45^{lo}B220^{+}CD19^{+}IgM^{+}IgD^{+/-}$ bone marrow cells from pups from control and *S. mansoni* infected mothers. Each group represents a pool of >4 mice. Transcriptional markers shown as FeaturePlots used to identify different clusters.

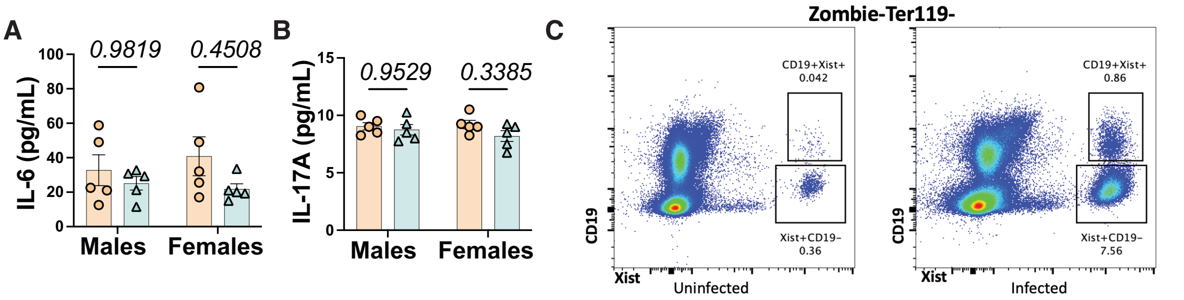


**Figure S8: Maternal schistosomiasis causes offspring immunity defects independent of known IL-6 pathway.** (**A**) Cytokine levels of IL-6 and (**B**) IL-17A from serum of pups from infected and control mothers. Statistics determined by two-way ANOVA. (**C**) *Xist* staining in peripheral blood mononuclear cells of uninfected and *S. mansoni* infected mothers by flow-fish.

88. D. Cortes-Selva, A. Ready, L. Gibbs, B. Rajwa, K. C. Fairfax, IL-4 promotes stromal cell expansion and is critical for development of a type-2, but not a type 1 immune response. *Eur J Immunol* **49**, 428-442 (2019).
89. M. Yousefi *et al.*, Comparative in vitro and in vivo assessment of toxin neutralization by anti-tetanus toxin monoclonal antibodies. *Hum Vaccin Immunother* **10**, 344-351 (2014).
90. A. Butler, P. Hoffman, P. Smibert, E. Papalexi, R. Satija, Integrating single-cell transcriptomic data across different conditions, technologies, and species. *Nat Biotechnol* **36**, 411-420 (2018).
91. Y. Hao *et al.*, Integrated analysis of multimodal single-cell data. *Cell* **184**, 3573-3587 e3529 (2021).
92. T. Stuart *et al.*, Comprehensive Integration of Single-Cell Data. *Cell* **177**, 1888-1902 e1821 (2019).
93. R. D. Lee *et al.*, Single-cell analysis identifies dynamic gene expression networks that govern B cell development and transformation. *Nat Commun* **12**, 6843 (2021).
94. N. Borcherding, N. L. Bormann, G. Kraus, scRepertoire: An R-based toolkit for single-cell immune receptor analysis. *F1000Res* **9**, 47 (2020).
95. J. Samir, S. Rizzetto, M. Gupta, F. Luciani, Exploring and analysing single cell multi-omics data with VDJView. *BMC Med Genomics* **13**, 29 (2020).
96. G. A. Challen, N. Boles, K. K. Lin, M. A. Goodell, Mouse hematopoietic stem cell identification and analysis. *Cytometry A* **75**, 14-24 (2009).
97. E. Sitnicka *et al.*, Key role of flt3 ligand in regulation of the common lymphoid progenitor but not in maintenance of the hematopoietic stem cell pool. *Immunity* **17**, 463-472 (2002).
98. C. Chu, J. Quinn, H. Y. Chang, Chromatin isolation by RNA purification (ChIRP). *J Vis Exp*, (2012).
99. C. Chu *et al.*, Systematic discovery of Xist RNA binding proteins. *Cell* **161**, 404-416 (2015).
100. G. R. Pullen, M. G. Fitzgerald, C. S. Hosking, Antibody avidity determination by ELISA using thiocyanate elution. *J Immunol Methods* **86**, 83-87 (1986).
